# Age-related reorganization of functional network architecture in semantic cognition

**DOI:** 10.1101/2022.05.10.491274

**Authors:** Sandra Martin, Kathleen A. Williams, Dorothee Saur, Gesa Hartwigsen

## Abstract

Semantic memory is a fundamental human ability which is central to communication. Although it is usually well preserved in healthy aging, memory problems in verbal communication due to slowed access and retrieval processes are a common complaint with increasing age. So far, the neural bases of this paradox remain poorly understood. The current neuroimaging study investigated age differences in the functional network architecture during semantic word retrieval in young and older adults. Using group spatial independent component analysis, we defined functional networks for verbal semantic fluency. Combining task-based functional connectivity, graph theory and cognitive measures of fluid and crystallized intelligence, our findings show age-accompanied large-scale network reorganization even when older adults have intact word retrieval abilities. In particular, functional networks of older adults were characterized by reduced decoupling between systems, reduced segregation and efficiency, and a larger number of hub regions relative to young adults. Exploring the predictive utility of these age-related changes in network topology revealed high, albeit less efficient, performance for older adults whose brain graphs showed stronger dedifferentiation and reduced distinctiveness. Our results extend theoretical accounts on neurocognitive aging by revealing the compensational potential of the commonly reported pattern of network dedifferentiation when older adults can rely on their prior knowledge for successful task processing. However, we also demonstrate the limitations of such compensatory reorganization processes and demonstrate that a youth-like network architecture in terms of balanced integration and segregation is associated with more economical processing.

**Significance Statement:** Cognitive aging is associated with widespread neural reorganization processes in the human brain. However, the behavioral impact of such reorganization is not well understood. Here, we used task-based fMRI to demonstrate a large-scale reorganization of brain networks in older adults even when their semantic abilities are intact. In particular, functional networks of older adults were characterized by increased coupling between different systems, reduced segregation and efficiency, and a larger number of hub regions relative to young adults. Associating these changes with behavior revealed high, albeit less efficient, performance for networks in older adults showing stronger dedifferentiation and reduced distinctiveness. Our results highlight the compensatory potential of network reconfiguration with age, but also reveal the limitations of such reorganization processes.

## Introduction

Semantic memory refers to the general knowledge of words, concepts, and ideas we accumulate across the lifespan. It is a fundamental human ability and central to communication. Unlike other cognitive domains, semantic memory is usually preserved through adulthood into very old age (Verhaegen et al., 2003), thus enabling communication abilities to remain largely intact in healthy aging. Nonetheless, memory problems in verbal communication, such as finding the right word and tip-of-the-tongue episodes, are a common complaint with increasing age (Burke and Shafto, 2004). This paradox has been explained in terms of less efficient access and retrieval processes during language production that rely on semantic and cognitive control functions like working memory, attention, and inhibitory control, and are well established to steadily decline with age (Hedden and Gabrieli, 2004). However, little is known about the neural mechanisms underlying those changes in access to semantic memory with age.

The field of network science provides tools to model and explore organization principles of complex systems such as the human brain (Rubinov and Sporns, 2010). Studies in young adults have revealed a topological organization of the brain that combines local information processing with global information integration aimed at optimizing global cost efficiency (“small world” organization; Bassett et al., 2009; Bullmore and Sporns, 2012). Age-related changes to this modular organization have been described as general decline of network segregation in the form of decreased within- and enhanced between-network connectivity (Chan et al., 2014; Setton et al., 2022). Moreover, increasing age has been associated with reduced small-world organization, modularity, and local and global efficiency of functional brain networks (Betzel et al., 2014; Geerligs et al., 2015; Chong et al., 2019). The impact of such reorganization on cognition remains debated. Most studies associated neural dedifferentiation with performance decline (Chan et al., 2014; Sala-Llonch et al., 2014; Chong et al., 2019), whereas some have pointed towards a pattern of compensational response (Stumme et al., 2020).

To date, most results stem from resting-state fMRI investigations or task-based studies in domains primarily affected by age, such as episodic and working memory. However, important insight can be gained by investigating domains which rely on semantic cognition like language and creativity. Here, older adults might benefit from increased connectivity between usually anticorrelated networks such as executive and default networks since they can depend on prior knowledge to maintain high performance (Spreng et al., 2016; Adnan et al., 2019). In this context, semantic fluency tasks are especially valuable since they tap into semantic memory but also cognitive control and are often linked to preserved albeit slower performance with age (Gordon et al., 2018). Previous studies revealed age-related reduced functional connectivity within domain-specific networks, however, without affecting behavioral performance (Marsolais et al., 2014; Ferré et al., 2020). Additionally, we recently showed that increased crosstalk between domain-general networks is essential for successful task processing, independent of age, when access to semantic memory is required (Martin et al., 2022). Thus, domains that are usually well-preserved in aging inform the current understanding of age-accompanied changes in functional brain networks and their behavioral relevance.

The present study contributes to this field by exploring age-related reorganization of functional networks during a semantic word retrieval task. Networks of task-based functional connectivity in groups of healthy young and older adults were derived via data-driven, multivariate methods. We were interested in age differences in the coupling of task-relevant networks and their behavioral relevance. Further, we applied graph-theoretical measures of brain system segregation, integration, and network hubs to investigate the network topology in young and older adults, and related these measures to participants’ in-scanner task performance and abilities of fluid and crystallized intelligence. Exploring task-based network topologies as a function of cognitive performance in a domain that is usually well preserved with age enabled us to gain key insights into age-related reorganization processes and to inform theoretical accounts regarding compensatory and detrimental effects of neurocognitive aging on behavior.

## Materials and Methods

### Participants

Participants consisted of 31 healthy older adults (*n* = 31, mean = 65.5, SD = 2.75, range = 60–69 years) and 30 healthy young adults (*n* = 30, mean = 27.6, SD = 4.3, range = 21–34 years), which is the same sample as described previously (Martin et al., 2022). Data of three older participants as well as single runs of six participants had to be excluded due to strong motion during fMRI (>1 voxel size), leading to a final sample size of 28 participants in the older group. While both groups were matched for gender, participants in the young group had significantly more years of education (*t*(55.86) = 5.21, p < 0.001). Inclusion criteria were native German speaker, right-handedness, normal hearing, normal or corrected-to-normal vision, no history of neurological or psychiatric conditions, and no contraindication to magnetic resonance imaging. Older adults were additionally screened for cognitive impairments with the Mini-Mental State Examination (Folstein et al., 1975); all ≥26) points and for depression with the Beck Depression Inventory (Beck et al., 1996); all ≤14 points). A battery of neuropsychological tests was administered probing semantic knowledge as well as verbal- and non-verbal executive functions (Table S1). Differences between age groups for neuropsychological measures were determined with two-sample t-tests. Consistent with previous research, older adults only performed better for the measure of semantic memory (spot-the-word test; *t*(54.39) = 3.14, *p* = 0.003), indicating a maintenance of semantic knowledge and an increase in vocabulary with age (Verhaegen et al., 2003), while young adults performed better on all other tests (all at *p* < 0.01), which is consistent with the assumption of a general age-related decline of executive functions (Hedden and Gabrieli, 2004). For all reported correlation analyses, neuropsychological measures were summarized via exploratory factor analysis. Results revealed an “executive functions” factor with high loadings on Trail Making Tests A (0.8) and B (0.71), Digit Symbol Substitution Test (0.73), and reading span test (0.45), and a “semantic memory” factor with spot-the-word test (0.5) and verbal fluency tests for hobbies (0.44) and surnames (0.98). Prior to the experiment, participants gave written informed consent. The study was approved by the local ethics committee of the University of Leipzig and conducted in accordance with the Declaration of Helsinki.

### Experimental design

The experimental procedure is reported in detail in previous work (Martin et al., 2022) and briefly summarized here. Participants completed one experimental session which consisted of two runs of the fMRI experiment and neuropsychological tests, and lasted two hours in total. Experimental tasks consisted of a paced overt semantic fluency task and a control task of paced overt counting, which were implemented in a block design in the scanner (Figure 1). For the semantic fluency task, participants were asked to produce exemplars for 20 semantic categories, which were divided in 10 easy (e.g., colors) and 10 difficult (e.g., insects) categories based on a separate pilot study in healthy young and older adults (Martin et al., 2022). Task blocks were 43 s long and separated by rest blocks of 16 s. Each block started with a 2 s visual word cue indicating whether participants were expected to generate category exemplars or count forward (1 to 9) or backward (9 to 1). This was followed by nine consecutive trials of the same category or counting task, respectively. Trials within one block were separated by inter-stimulus intervals of 2-4 s. Participants were instructed to generate one exemplar for a category or one number per trial, which was indicated by a green cross on the screen, and to pause when the cross turned red. They were told not to repeat items and to say “next” if they could not think of an exemplar for the respective category. Each run contained 10 semantic fluency blocks, divided in easy and difficult categories, and 10 counting blocks, consisting of forward and backward counting, thus resulting in a total duration of 19.4 min per run. The order of blocks was counter-balanced and pseudo-randomized across participants. Before the fMRI experiment, participants received instructions and practiced the task with a separate set of categories outside the scanner. Stimuli were presented using the software Presentation (Neurobehavioral Systems, Berkeley, USA; version 18.0). Answers were recorded via a FOMRI III microphone (Optoacoustics, Yehuda, Israel). After the experiment, response recordings were analyzed for verbal answers and onset times after being cleaned from scanner noise via Audacity software (version 2.3.2) and transcribed by three independent raters.

**Figure 1.**
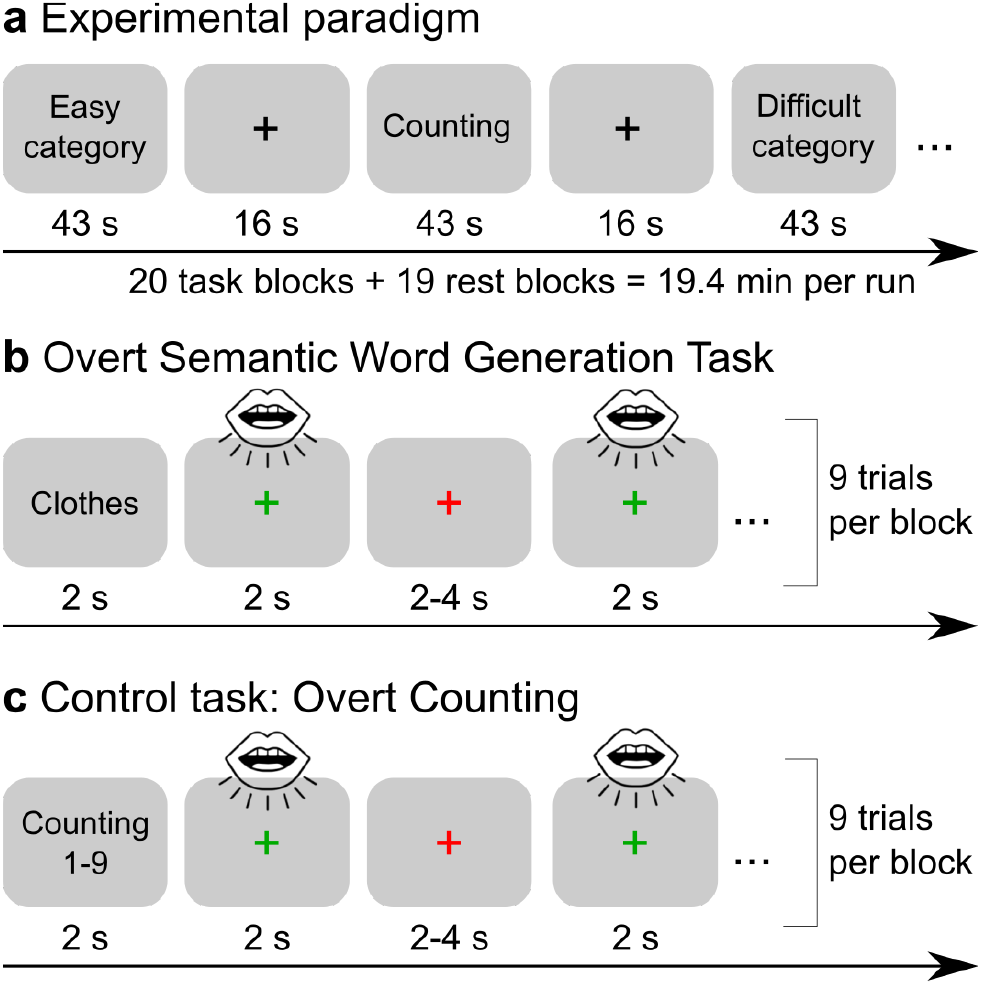
Experimental design. (a) The fMRI experiment consisted of 43-s task blocks of overt paced semantic fluency and counting, which were presented in a pseudorandomized order and separated by 16-s rest periods. An example for each task is shown in (b) and (c). There were 10 blocks per run for each task. At the beginning of a task block, a 2 s visual word cue indicated whether participants were expected to generate category exemplars or count forward (1 to 9) or backward (9 to 1). Participants were instructed to produce exactly one exemplar for a category or to say one number when the fixation cross turned green and to pause when the cross turned red. If they could not think of an exemplar, they were instructed to say “next”. Each task block contained 9 trials of the same semantic category/counting task which were separated by jittered inter-stimulus intervals.

### fMRI data acquisition and preprocessing

fMRI data were collected on a 3 T Prisma scanner (Siemens, Erlangen, Germany) with a 32-channel head coil. For the acquisition of functional images, a multiband dual gradient-echo echo-planar imaging sequence was used for optimal blood oxygenation level-dependent (BOLD) sensitivity throughout the entire brain (Poser et al., 2006; Halai et al., 2014). The following scanning parameters were applied: TR = 2000 ms; TE = 12 ms, 33 ms; flip angle = 90°; voxel size = 2.5 × 2.5 × 2.75 mm with an inter-slice gap of 0.25 mm; FOV = 204 mm; multiband acceleration factor = 2. To increase coverage of anterior temporal lobe (ATL) regions, slices were tilted by 10° of the AC-PC line. 616 images consisting of 60 axial slices in interleaved order covering the whole brain were continuously acquired per run. Additionally, field maps were obtained for later distortion correction (TR = 8000 ms; TE = 50 ms). This study analyzed the data from echo 2 (TE = 33 ms) since preprocessing was performed using the software fMRIPrep (Esteban et al., 2019), which currently does not support the combination of images acquired at different echo times. We chose to use results from preprocessing with fMRIPrep since this pipeline provides state-of-the-art data processing while allowing for full transparency and reproducibility of the applied methods and a comprehensive quality assessment of each processing step that facilitates the identification of potential outliers. We also double-checked results from preprocessing with fMRIPrep with a conventional SPM preprocessing pipeline of both echoes. The comparison of both pipelines did not reveal big differences in analysis results. A high-resolution, T1-weighted 3D volume was obtained from our in-house database (if it was not older than two years) or collected after the functional scans using an MPRAGE sequence (176 slices in sagittal orientation; TR = 2300 ms; TE = 2.98 ms; flip angle = 9°; voxel size = 1 × 1 × 1 mm; no slice gap; FOV = 256 mm). Preprocessing was performed using fMRIPprep 20.2.3 (Esteban et al., 2019) which is based on Nipype 1.6.1 (Gorgolewski et al., 2011). In short, preprocessing steps included skull stripping, distortion correction, co-registration, slice timing correction, and calculation of several confounding time-series for each of the two BOLD runs per participant. Anatomical T1-weighted images were skull-stripped, segmented, and spatially normalized. For spatial normalization to standard space, the Montreal Neurological Institute (MNI) ICBM 152 non-linear 6th Generation Asymmetric Average Brain Stereotaxic Registration Model (MNI152NLin6Asym) was entered as output space in fMRIPrep. For more details on the preprocessing pipeline, see the section corresponding to workflows in fMRIPrep’s documentation (https://fmriprep.org/en/20.2.3/workflows.html). After preprocessing, 29 volumes from the beginning of each run were discarded since they were collected for the combination of the short and long TE images. This yielded 587 normalized images per run which were included in further analyses.

### Independent component analysis

We applied group independent component analysis (ICA) to define spatially independent task-active networks in a data-driven manner. ICA has been shown to decompose fMRI time series into reliable functionally connected components with the advantage of simultaneously removing non-neural fluctuations through the identification of artefactual components (Griffanti et al., 2014).

Preprocessed, normalized data were smoothed with a 5 mm^3^ FWHM Gaussian kernel and entered into a general linear model for each participant and session using Statistical Parametrical Mapping software (SPM12; Wellcome Trust Centre for Neuroimaging), implemented in MATLAB (version 9.10/R2021a). GLMs included regressors for the task blocks (semantic fluency and counting) as well as nuisance regressors consisting of the six motion parameters and individual regressors for strong volume-to-volume movement as indicated by values of framewise displacement > 0.9 (Siegel et al., 2014). Additionally, an individual regressor of no interest was included in the design matrix if a participant had missed a whole task block during the experiment (*n* = 10). Before model estimation, a high-pass filter with a cut-off at 128 s was applied to the data.

Preprocessed, normalized and smoothed data were analyzed using the Group ICA of fMRI Toolbox (GIFT v4.0c). Dimensions were reduced to 55 using minimum description length information criteria. Icasso was repeated 50 times to ensure reliability of the decomposition, and group-level ICs were back-reconstructed to the participant level using the group-information guided ICA (GICA3) algorithm (Calhoun et al., 2001). We calculated group ICA treating all participants as one group to ensure that the same components were identified in both groups. We discarded those components related to banding artifacts and noise after careful visual inspection of the spatial maps according to established criteria (Griffanti et al., 2014); see Figure S1 for an overview of all 55 ICs). From the resulting 13 non-noise components, low-level sensory components including auditory, sensorimotor, and visual networks were identified and removed since their roles were beyond the scope of our investigation. To characterize the spatial extent of the seven remaining components at the group level, we calculated one-sided t-tests for participants’ spatial maps. A gray matter mask that restricted statistical tests to voxels in the cerebrum was applied to all group-level analyses. Results were corrected for multiple comparisons using a peak level threshold at *p* < 0.05 with the family-wise error (FWE) method and a cluster-extent threshold of 10 voxels.

### Brain network construction

Brain networks were constructed based on the seven selected component maps of the ICA. To determine network labeling of the thresholded maps, we used the Jaccard index (*J*), a measure of spatial similarity (Jaccard, 1912). By calculating the ratio of overlapping voxels in two binary spatial network maps relative to all active voxels in either image, the Jaccard index can be used as a measure to assess the fit between a spatial component map (*A*) and a template image (*B*):

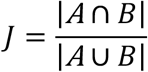

The index ranges from 0 to 1, with a high Jaccard index denoting high similarity of two spatial maps. It has been used previously to assess similarity of brain activation maps with template network parcellations (Jackson et al., 2019; Gordon et al., 2020). We defined a minimum threshold of J = 0.15 to consider a network template for a spatial component mask (Jackson et al., 2019). Next, if two components were best described by the same network template thereby indicating that the network might have split up in multiple components, we assessed the similarity of the combined component maps to the template. If the combined map reached a higher similarity index than each component individually, the combination was kept as a reflection of the respective network.

As template masks, we used the 17-networks functional connectivity-based parcellation scheme (Yeo et al., 2011) as well as the network masks of general semantic cognition and semantic control defined in a meta-analysis (Jackson, 2021). We included separate template masks for semantic cognition in our analysis to account for the semantic nature of our task. We also probed similarity of Jaccard indices with a 7-networks parcellation scheme (Yeo et al., 2011). While the results for the 7-networks parcellation generally agreed with the more fine-grained parcellation, the 7-networks parcellation resulted in three components showing high spatial similarity with the default network template. However, differential roles have been reported for subsystems of the default network when access to semantic memory is required (Smallwood et al., 2021). Specifically, the dorsal medial subsystem of the default network (“Default B” in the 17-networks parcellation scheme) has been shown to broadly overlap with a left-lateralized temporal-frontal semantic network (Lambon Ralph et al., 2017; Smallwood et al., 2021). Since we were interested in the age-dependent interplay of domain-specific and domain-general networks in semantic cognition, the remaining analyses were based on the 17-networks parcellation scheme.

Based on the results of the Jaccard index, each thresholded component map was inclusively masked by the respective resampled template network. We were interested in the effect of age on the functional connectivity within and between selected networks. In a first step, to explore functional connectivity between networks, we extracted averaged time series across all voxels within one masked component, thus leading to seven time series per participant and run. Second, networks were further parcellated into distinct regions of interest (ROIs) based on peak maxima of activated clusters. ROIs were created for all peak maxima of a significant cluster (up to three ROIs per cluster) using the MarsBar toolbox (Brett et al., 2002). To this end, identified clusters were extracted from the thresholded and masked component maps, spheres of 5 mm surrounding each maximum coordinate were created, and, in a last step, both images were combined. In this way, we ensured that ROIs would only contain voxels that were included in the group-level statistics. Parcellating the seven network components based on strongest correlation peaks led to 126 cortical ROIs per participant and run.

Functional time series were extracted for the seven ROIs and 126 ROIs parcellation schemes from non-smoothed functional data. To account for motion artifacts and other signal confounds, the following denoising pipeline was applied during time series extraction: 24 realignment parameters (six motion parameters, temporal derivatives, and quadratic terms), global signal, and top five aCompCor components for white matter and cerebral spinal fluid, respectively. Censoring included a framewise displacement threshold of 0.9 mm and 18 discrete cosine-basis regressors to account for signal drifts. All these regressors were combined in a design matrix and removed from the data in a single step (Hallquist et al., 2013; Lindquist et al., 2019). The denoising strategy was based on recent recommendations (Mascali et al., 2021) that compared the performance of different denoising pipelines for analysis of task-based functional connectivity. Consistent with previous research on resting-state functional connectivity (Ciric et al., 2017; Parkes et al., 2018), the authors reported that the inclusion of global signal in a denoising pipeline markedly reduced global motion artifacts and led to more comparable results across conditions in task-based functional connectivity data (Mascali et al., 2021). Further, time series were detrended and demeaned, and functional images were masked with a subject-specific, resampled gray matter mask before denoising. During signal extraction for the set of 126 ROIs, the number of voxels per ROI and participant were extracted. ROIs for which more than 15% of participants did not show any signal coverage were excluded. The resulting 121 ROIs were used for the remaining analyses.

### Functional connectivity matrices

We applied correlational psychophysiological interaction analyses (cPPI; Fornito et al., 2012) to obtain connectivity terms that describe task-related interactions between our networks and regions of interest. In contrast to traditional PPI analyses, cPPI results in undirected, symmetrical connectivity matrices that are based on pairwise partial correlations between ROIs. We calculated cPPI for our contrast of interest semantic fluency > counting, separately for the 7-networks and 121-ROIs parcellations. In brief, the deconvolved time series for each ROI was multiplied with the task time course from the first-level GLM design matrix and convolved with a canonical HRF to form a PPI term. Pairwise partial correlations were estimated between PPI terms of two regions while controlling for the observed BOLD signal in both regions, the original task regressor and average in-scanner head motion (mean FD). Connectivity matrices were calculated for each run separately and then averaged, resulting in a 7 × 7 and 121 × 121 correlation matrix per participant. Subsequently, correlation coefficients were Fisher-transformed to z values.

### Network measures

#### Within- and between-network functional connectivity

Within- and between-network functional connectivity were explored for the 7-networks and 121-ROIs connectivity matrices in both age groups. Using the connectivity matrices with seven networks allowed us to investigate the coupling and decoupling between task-relevant networks while the more fine-grained parcellation provided additional insights into the coupling of regions within distinct networks. All subsequent network measures were based on the 121-ROIs connectivity matrices.

#### Brain system segregation

We calculated global segregation as previously implemented by Chan and colleagues (Chan et al., 2014, 2021; Wig, 2017), using the unthresholded, weighted connectivity matrices. In line with previous work on functional connectivity in healthy aging (Chong et al., 2019; Chan et al., 2021), we excluded negative correlations from segregation and integration analyses by setting them to zero. Excluding negative correlations has been shown to improve the reliability of graph measures (Wang et al., 2011) and to help avoid interpretational difficulty, for example when it comes to concepts like shortest paths (Fornito et al., 2016). Building upon the network parcellation of our ICA analysis, each functional network was treated as a distinct system, and segregation was computed as the difference between mean within-system 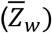 and mean between-system 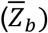 correlations divided by mean within-system correlation as shown in the following equation:

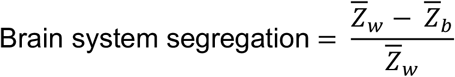

A higher ratio score denotes greater separation of functional systems.

We also calculated segregation values for each functional network individually such that within-system connectivity 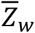 represents the mean of all edges (correlations) between pairwise nodes that belong to the same network and between-system connectivity 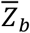 reflects the mean of all edges between nodes of the respective network and all other nodes.

#### Edge filtering

Most graph theoretical measures require some form of filtering to obtain a sparse graph that is more likely to represent true functional connectivity than a maximally dense graph as produced by a correlation matrix (Fornito et al., 2016). While threshold-based filtering methods like proportional or absolute thresholding are commonly applied in network neuroscience, they are driven by an arbitrary choice of the respective threshold and suffer from low reliability (Luppi and Stamatakis, 2021). To avoid these pitfalls and based on recent research on the reliability of graph construction in neuroscience (Jiang et al., 2021; Luppi and Stamatakis, 2021), we calculated the orthogonalized minimum spanning tree (OMST; Dimitriadis et al., 2017) on the weighted functional connectivity matrices. Apart from its high reliability, the OMST has several advantages compared to commonly applied threshold-based methods of graph construction: It adheres to the intrinsic topological structure of the brain network by resulting in a fully connected, weighted graph and offers a data-driven method of individualized network construction accounting for each individual’s optimal state of economic wiring in terms of cost and efficiency. In contrast to the original minimum spanning tree (MST), the OMST filters connectivity networks until the highest global cost efficiency (GCE) of a graph is reached while including both strong and weak connections and preserving the same mean degree across groups.

The OMST was calculated in three steps as described by Dimitriadis et al. (2017): (a) the MST of a graph is defined; (b) the corresponding edges of the MST are removed from the original graph by setting edge weights to 0; (c) steps (a) and (b) are repeated until the GCE of the graph is optimized. GCE is defined as the global efficiency minus cost, where cost corresponds to the total weights of the selected edges of the OMST divided by the sum of the edges of the original fully weighted graph (Bassett et al., 2009). The final OMST is constructed by combining all the removed, non-overlapping MSTs. To show that the OMST indeed results in higher GCE than other filtering methods, we compared the GCE for OMST, MST, and a method of proportional thresholding where we used a common range of 5-20% strongest edge weights of a graph (Figure S3). To avoid differences in graph measures caused by the number of nodes in a graph, we excluded all nodes where at least one participant had no signal during construction of matrices. This resulted in a 104×104 matrix per participant, which was used for construction of OMST and all subsequent measures.

#### Brain system integration

We calculated global efficiency as a measure of system-wide integration. It is defined as the average of the inverse shortest path length between all pairs of nodes in a graph and is thus a measure of efficient signal transmission (Latora and Marchiori, 2001; Rubinov and Sporns, 2010).

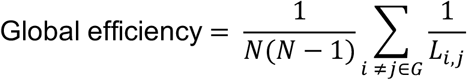

Global efficiency was based on the individual OMSTs using the reciprocal edge weights to obtain a distance matrix where high weights signify short paths between nodes.

#### Global network hubs

We identified hubs via the normalized participation coefficient (PC; Pedersen et al., 2020). The PC provides insight into the functional role of a node. Specifically, it evaluates whether a node mainly interacts with nodes from its community or multiple communities of a network (Guimerà and Nunes Amaral, 2005). In network neuroscience, PC has been applied to define nodes that are important for communication between communities (connector hubs) and nodes that are central to the communication within communities (provincial hubs; Cohen and D’Esposito, 2016; Bertolero et al., 2017, 2018). Recently, it has been shown that the conventional measure of PC is strongly influenced by the size and connectedness of its community leading to a reduced interpretational value of this graph measure (Pedersen et al., 2020). Thus, a normalized version of the PC has been introduced that accounts for these differences in real-world networks while preserving its meaning though the comparison with null models. It is calculated similarly to the original PC as one minus the ratio between the degree *k* of node *i*. with nodes in its community *m* and the degree of node *i*. with all other nodes in the network. However, a normalization factor is added by subtracting the median degree of this node in a series of random networks:

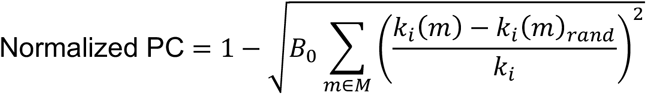

We calculated 100 random networks for each node. Connector hubs were then defined as nodes with a PC value of at least 1SD above the mean in each age group.

### Statistical analyses

#### Age-related changes for within- and between-network functional connectivity

To assess differences between age groups for within- and between-network connectivity, we ran two-sample t-tests for each edge of the 7-network and 121-ROIs connectivity matrices within the Network-Based Statistics toolbox (NBS; Zalesky et al., 2010). NBS applies cluster-based thresholding to correct for multiple comparisons using permutation testing. In contrast to more conventional procedures for controlling the family-wise error rate, such as the false discovery rate, NBS considers connected components in networks (graphs), which makes it especially suited for network statistics. We set an initial cluster-forming threshold at *p* < 0.01 (two-sided test; *t* = 2.67) and an FWE-corrected significance threshold at *p* < 0.05 with 10,000 permutations. To control for the influence of motion on functional connectivity, the average in-scanner head motion per participant was included as a covariate. Average head motion was defined as the mean FD based on the calculation of the root mean square deviation of the relative transformation matrices (Jenkinson et al., 2002).

#### Age-related changes for network measures of segregation and integration

Linear mixed-effects models were set up to examine how the dependent variables brain system segregation, individual network segregation, global efficiency, and nodal participation coefficient were predicted by age group. We included in-scanner head motion (mean framewise displacement) as covariate and a random intercept for participants. Models were calculated as follows:

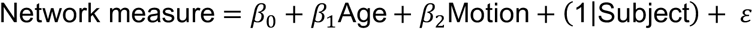

Significance values were obtained by likelihood ratio tests of the full model with the effect in question against the model without the effect in question.

#### Association between network measures and cognitive performance

For those network measures that showed differences between young and older adults, we further examined their association with participants’ cognitive performance for the in-scanner task and the neuropsychological test battery. Analyses were performed using mixed-effects models with a logistic regression for accuracy data due to their binomial nature and a linear regression for log-transformed response time data. We only analyzed response times for correct reactions for the semantic fluency task since our connectivity values were also based on our contrast of interest semantic fluency > counting. Models contained fixed effects for the respective mean-centered network measure (between-network functional connectivity, brain system segregation, individual network segregation, and global efficiency) and age group as well as their interaction term, and random intercepts for participants and semantic categories. Further, mean-centered values of mean FD and education were entered as covariates. Models were set up as shown in the following equation:

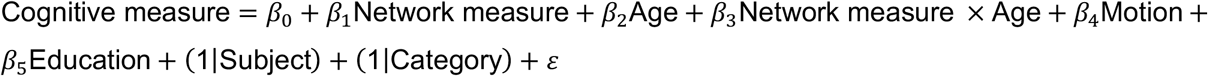

where cognitive measure denotes accuracy and response time, respectively. Significance values were obtained via likelihood ratio tests. We applied sum coding (ANOVA-style encoding) for all categorical predictors. Results were corrected for multiple comparisons using the Bonferroni-Holm method.

We performed correlation analyses with the neuropsychological tests that had been assessed outside of the scanner. Due to the collinearity of some neuropsychological tests, we ran an exploratory factor analysis on the standardized test scores using maximum likelihood estimation and varimax rotation. Based on the hypothesis test (*χ*^2^ = 14.04, *p* = 0.081), two factors with an eigenvalue > 1 were chosen. For subsequent correlations with network measures, participant factor scores extracted via regression methods were used.

All statistical models except for NBS were performed using R 4.1.0 via RStudio (R Core Team, 2021) and the package lme4 (Bates et al., 2015). Results were visualized using the ggplot2 (Wickham, 2016) and ggeffects (Lüdecke, 2018) packages. If applicable, post-hoc comparisons were applied using the package emmeans (Lenth, 2020). The exploratory factor analysis was calculated with the stats package (R Core Team, 2021). OMSTs and all graph theory measures were calculated in Matlab using the Brain Connectivity toolbox (Rubinov and Sporns, 2010) and publicly available scripts for OMST and normalized PC. Chord diagrams were generated with the circlize package (Gu et al., 2014), spring-embedded plots using the igraph package (Csardi and Nepusz, 2006), and force-directed plots using the ForceAtlas2 algorithm for R available on Github (https://github.com/analyxcompany/ForceAtlas2).

### Data and code availability

All behavioral data and raw data of functional connectivity and graph-theoretical measures are available in a public repository at https://gitlab.gwdg.de/functionalconnectivityaging/mdn_lang_networkAnalysis. This repository also holds all self-written code for analyses and figures for this project. Raw neuroimaging data are protected under the General Data Protection Regulation (EU) and can only be made available from the authors upon reasonable request.

## Results

The main objective of this study was to investigate age-related changes in the functional network architecture engaged during the goal-directed access to semantic memory. For the in-scanner tasks of overt semantic fluency and counting, we fitted mixed-effects models accounting for individual variance of participants and semantic categories via random effects and the difference in years of education via covariate(Table S2). Likelihood-ratio tests showed that both age groups performed similarly (*x*^2^= 2.18, *p* = 0.14) and generally better for counting than semantic fluency (*x*^2^= 8.06, *p* = 0.005; Figure 2a). For response time, results showed an interaction between task and age group (*x*^2^= 79.73, *p* < 0.001) with older adults performing slower than young adults during the semantic fluency but not the counting task (Figure 2b).

**Figure 2.**
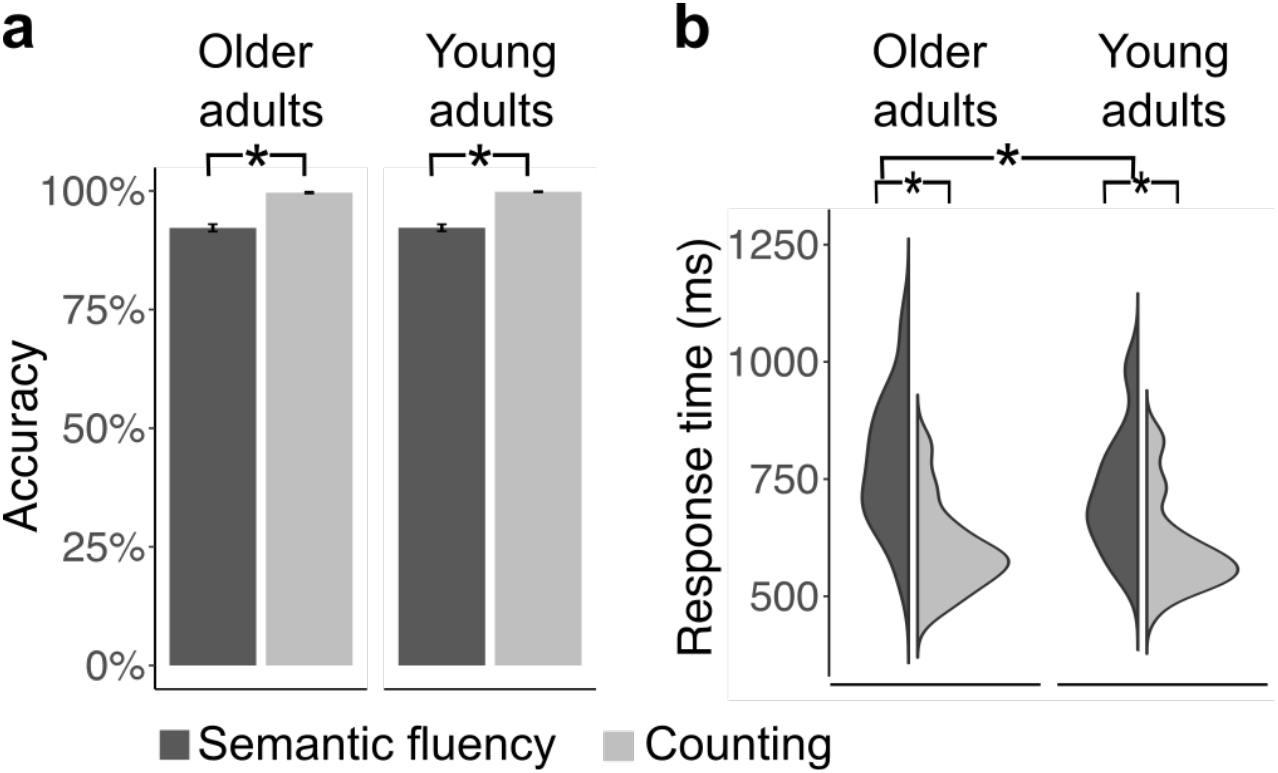
Behavioral results. Both groups performed better for counting than semantic fluency. While there was no effect of age for accuracy in either task (a), older adults performed slower than young adults during semantic fluency but not during counting (b). * p < 0.05.

### Goal-directed access to semantic memory involves default, semantic, and executive control networks

Using the data-driven method of group spatial independent component analysis (ICA) on the whole data set, we defined functional cortical networks for the semantic task. We identified seven components of interest which were submitted to one-sample t-tests and thresholded controlling the family-wise error (FWE) rate at peak level with *p* < 0.05 and a cluster-extent threshold of 10 voxels. Figure 3 shows the thresholded maps with their original component number. To determine which cognitive network best described each component, we calculated the Jaccard similarity coefficient (*J*) between our thresholded, binarized components of interest and template masks of common resting-state (Yeo et al., 2011) and semantic cognition networks (Jackson, 2021). Results showed similarity above threshold (*J* = 0.15) for all component maps with distinct cognitive networks (Table 1). For IC06, we found overlap with the frontoparietal control network C (CONT-C) and default mode network A (DMN-A). Although spatial similarity was marginally higher for CONT-C than DMN-A (*J*_Control C_ – *J*_Default A_ = 0.01), we refer to this component as part of the default system. Significant clusters included classic midline structures of the core default network (Smallwood et al., 2021) like posterior cingulate cortex, precuneus, and prefrontal cortex (Figure 3). An additional analysis of similarity coefficients between the component maps and the 7-networks parcellation (Yeo et al., 2011) revealed a stronger similarity with the default network as a whole for this component (*J*_Control_ – *J*_Default_ =-0.03; see Table S3 for results with the 7-networks parcellation). Furthermore, a second component (IC13) showed strong similarity with DMN-A. As described in Methods, we combined the component maps of IC06 and IC13 to assess whether this would lead to a numerical improvement of *J*. This was not the case with *J* = 0.21 for the combined components which was below the similarity coefficient of IC13 alone (*J* = 0.26). Thus, both components represented distinct parts of DMN-A and were hence included in subsequent analyses. For IC13, we further included default mode network C (DMN-C), which showed the second strongest overlap and was represented by significant clusters in bilateral parahippocampal gyri. Indeed, a combined template of DMN-A and DMN-C led to a numerical improvement in similarity compared to DMN-A alone (*J*_Default A + C_ – *J*_Defautt A_ = 0.091). Thus, to gain a comprehensive representation of the default network, both subsystems were combined and are referred to as default mode network A+C (DMN-A+C).

**Figure 3.**
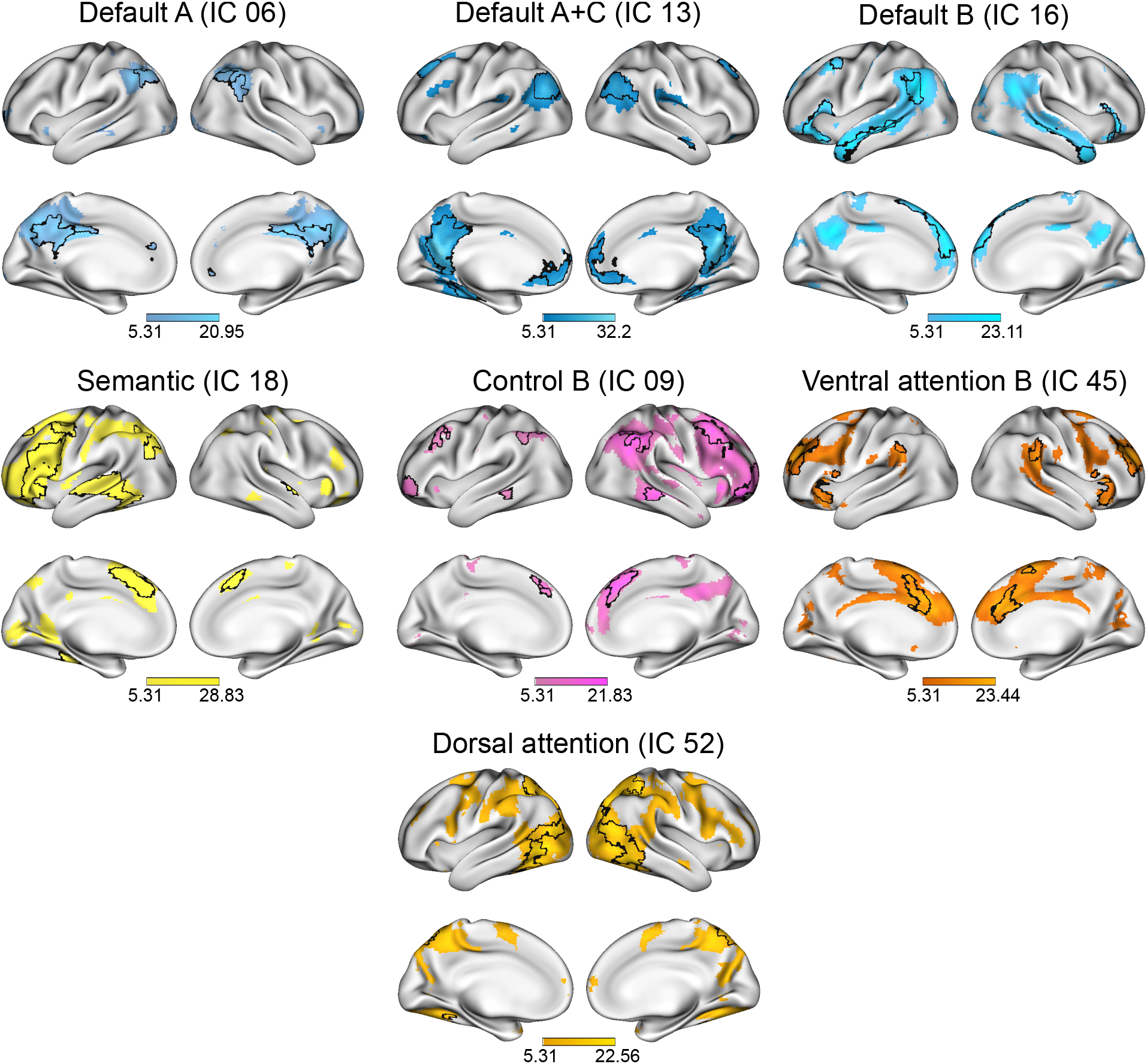
ICA-derived networks and their overlap with cognitive networks. T-scores from onesided t-tests (FWE-corrected *p* < 0.05 at peak level) are displayed for the seven selected component maps with their respective network label according to spatial similarity analysis. Overlaps between the thresholded component map and the spatially most similar cognitive network according to the Jaccard index are outlined on the surface of the brain. The areas of overlap were used for subsequent network analyses.

**Table 1.** Jaccard indices for independent components and cognitive networks

|  | IC06 | IC09 | IC13 | IC16 | IC18 | IC45 | IC52 |
| --- | --- | --- | --- | --- | --- | --- | --- |
| Frontoparietal control A | 0.054 | 0.133 | 0.032 | 0.013 | 0.151 | 0.083 | 0.109 |
| Frontoparietal control B | 0.091 | **0.210** | 0.028 | 0.073 | 0.125 | 0.050 | 0.018 |
| Frontoparietal control C | 0.168 | 0.020 | 0.066 | 0.010 | 0.010 | 0.028 | 0.044 |
| Default A | **0.154** | 0.089 | **0.255** | 0.149 | 0.040 | 0.054 | 0.019 |
| Default B | 0.039 | 0.069 | 0.031 | **0.263** | 0.082 | 0.098 | 0.010 |
| Default C | 0.014 | 0.010 | **0.122** | 0.008 | 0.026 | 0.001 | 0.020 |
| Dorsal attention A | 0.051 | 0.041 | 0.062 | 0.015 | 0.054 | 0.003 | **0.180** |
| Dorsal attention B | 0.008 | 0.038 | 0.006 | 0.006 | 0.071 | 0.053 | 0.123 |
| Limbic A | 0.000 | 0.001 | 0.002 | 0.014 | 0.002 | 0.002 | 0.011 |
| Limbic B | 0.001 | 0.007 | 0.015 | 0.004 | 0.009 | 0.005 | 0.002 |
| Ventral attention A | 0.042 | 0.012 | 0.023 | 0.015 | 0.033 | 0.124 | 0.065 |
| Ventral attention B | 0.014 | 0.074 | 0.001 | 0.031 | 0.059 | **0.195** | 0.039 |
| Somatomotor A | 0.022 | 0.052 | 0.000 | 0.039 | 0.036 | 0.029 | 0.028 |
| Somatomotor B | 0.000 | 0.016 | 0.038 | 0.009 | 0.033 | 0.011 | 0.015 |
| Temporal parietal | 0.001 | 0.029 | 0.014 | 0.118 | 0.023 | 0.035 | 0.034 |
| Central visual | 0.022 | 0.006 | 0.006 | 0.038 | 0.011 | 0.004 | 0.123 |
| Peripheral visual | 0.025 | 0.009 | 0.074 | 0.020 | 0.038 | 0.034 | 0.037 |
| General semantic cognition | 0.032 | 0.030 | 0.072 | *0.194* | **0.201** | 0.092 | 0.050 |
| Semantic control | 0.012 | 0.036 | 0.012 | 0.067 | *0.153* | 0.091 | 0.027 |

*Note.* The selected network labels for the respective independent components are shown in bold while all cognitive networks that showed a higher similarity coefficient than $J = 0.15$ are shown in italics.

Results of Jaccard calculations further revealed the following networks for the other components: default mode network B (DMN-B; IC16) with peak activations in bilateral middle temporal gyri (MTG), inferior and superior frontal gyri (IFG, SFG), and left angular gyrus (AG); semantic network (SEM; IC18) with strong overlap with the semantic control network and peak activations in left IFG, SFG, paracingulate gyrus, posterior superior temporal gyrus (STG), and AG; frontoparietal control network B (CONT-B; IC09) with large clusters in bilateral SFG and middle frontal gyri (MFG), AG, and posterior MTG; ventral attention network B (VAN-B; IC45) with peak activation in prefrontal cortex including paracingulate gyrus, bilateral IFG and supramarginal gyri; and dorsal attention network A (DAN-A; IC52) with large clusters in bilateral AG, and temporooccipital cortex. Statistical tables with all significant clusters are reported in Table S5.

### Stronger coupling of default and executive systems predicts intact but less efficient semantic retrieval in older adults

Graphs of task-related connectivity were derived via cPPI for matrices with seven and 121 nodes, respectively. We tested for statistically significant coupling differences between age groups by means of network-based statistics using permutation testing while controlling for in-scanner head motion (Figure 4a). Overall, the network of older adults showed reduced decoupling compared to young adults. Significantly stronger positive coupling was found in the graphs of older adults for the networks SEM with VAN-B, CONT-B with VAN-B, and DAN-A with VAN-B. A similar picture of age-related differences emerged for the more fine-grained graphs containing 121 nodes. Here, young adults generally showed stronger positive coupling within individual networks and between subsystems of the default network and stronger decoupling between different networks (Figure S2).

**Figure 4.**
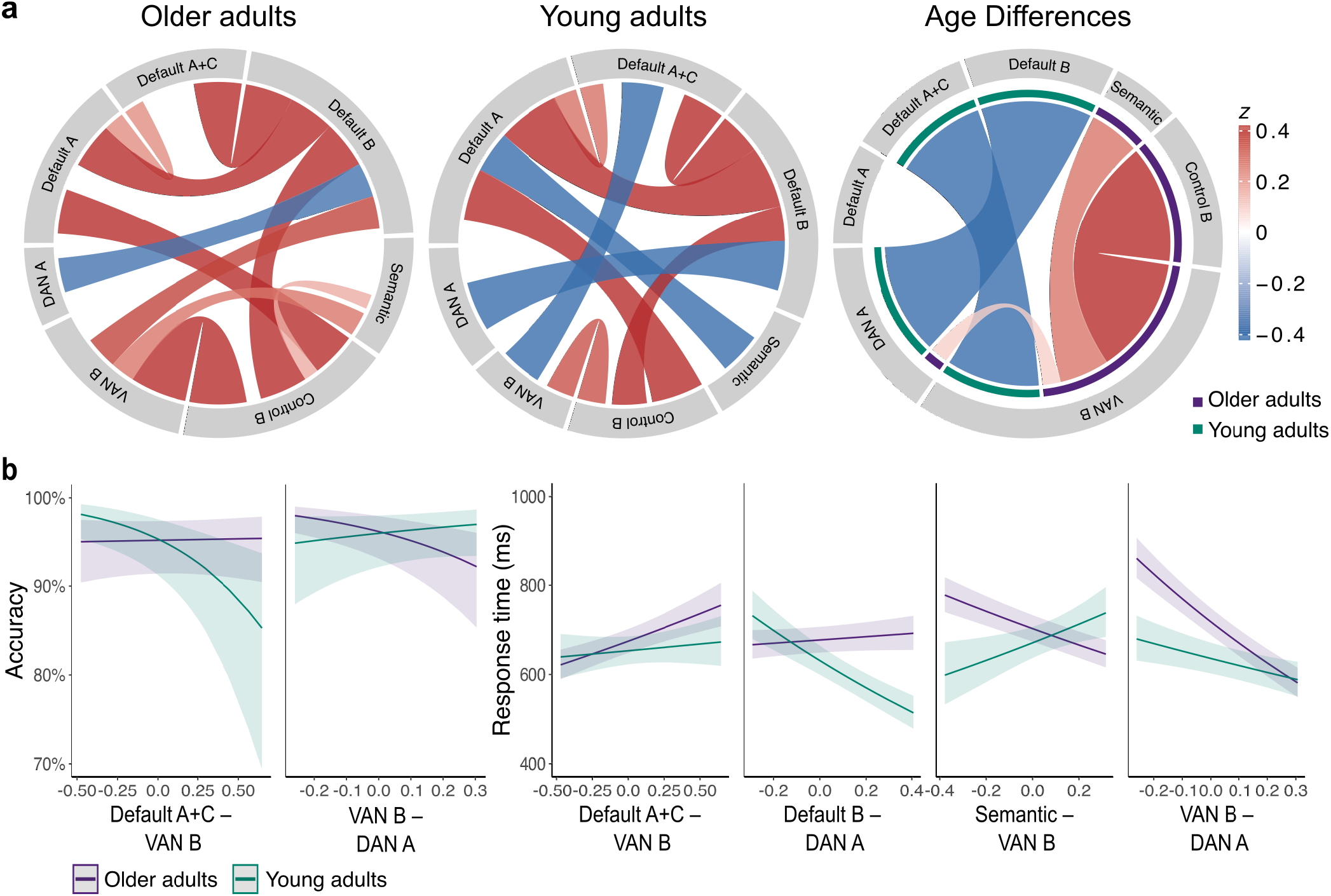
Functional coupling between task-relevant networks in young and older adults and their behavioral relevance. (a) Chord diagrams display significant results of functional coupling between ICA-derived networks. Connectivity values are partial correlations. The color intensity and width of a connection indicate its correlational strength. Red indicates coupling and blue indicates decoupling between networks. Chord diagrams of each age group are based on cPPI-derived significance values while age differences were assessed using permutation testing in networkbased statistics (cluster-forming threshold *p* = 0.01, FWE-corrected significance threshold *p* = 0.05 with 10,000 permutations). (b) Network connections that showed significant age differences were probed for their behavioral relevance. Plots show significant two-way interactions between age and the respective network pair for accuracy and response time data. Connectivity values were mean-centered for interaction analyses. Results were corrected for multiple comparisons using the Bonferroni-Holm method at *p* = 0.05. VAN ventral attention network, DAN dorsal attention network.

We probed the behavioral relevance of the network connection pairs that showed significant age differences by calculating mixed-effects models for accuracy and response time data (Figure 4b; Table S5-6). For accuracy, we found significant interactions between age and between-network connectivity for VAN-B with DMN-A+C (*x*^2^= 12.39, *p* = 0.002) and DAN-A (*x*^2^= 14.18, *p* < 0.001). Predicting response time revealed significant interactions between age and between-network connectivity for VAN-B with DMN-A+C (*x*^2^= 5.65, *p* = 0.035), SEM (*x*^2^= 25.75, *p* < 0.001), and DANA (*x*^2^= 28.81, *p* < 0.001), and for DAN-A with DMN-B (*x*^2^= 51.76, *p* < 0.001). For older adults, increased coupling between default and attention networks predicted high but less efficient performance, while increased coupling of SEM and VAN-B and between both attention systems (DAN-A and VAN-B) was associated with faster responses. A different picture emerged in young adults, where stronger coupling between default and executive systems predicted faster but poorer performance while increased connectivity between DAN-A and VAN-B was associated with better and faster reactions. These results suggest that both age groups showed distinct connectivity profiles, with older adults generally profiting from increased coupling between different cognitive systems and the opposite pattern for young adults.

### Reduced segregation and higher integration of task-relevant networks is associated with better and more efficient performance in older adults

Next, we investigated brain system segregation and integration to get a better understanding of age-related differences in whole-brain dynamics (Figure 5a). Segregation quantifies the presence of densely connected regions that form distinct subnetworks or communities in a global brain network (Figure 5b). Results of a linear mixed-effects model for global brain system segregation revealed a significant effect of age (*x*^2^= 11.23, *p* < 0.001) with young adults exhibiting stronger segregation than older adults (Figure 5c; Table S7). Examining the predictive value of segregation for in-scanner performance and neuropsychological measures revealed significant interactions between age and segregation for accuracy (*x*^2^= 9.54, *p* = 0.002), response time (*x*^2^= 71.15, *p* < 0.001), and a significant correlation of segregation with executive functions in young adults (*r* = 0.45, *p* = 0.013). For all interactions, increasing levels of segregation predicted better and faster performance in young adults. In contrast, increasing brain-wide segregation had no effect on accuracy but predicted faster responses in older adults (Figure 5c; Table S8).

**Figure 5.**
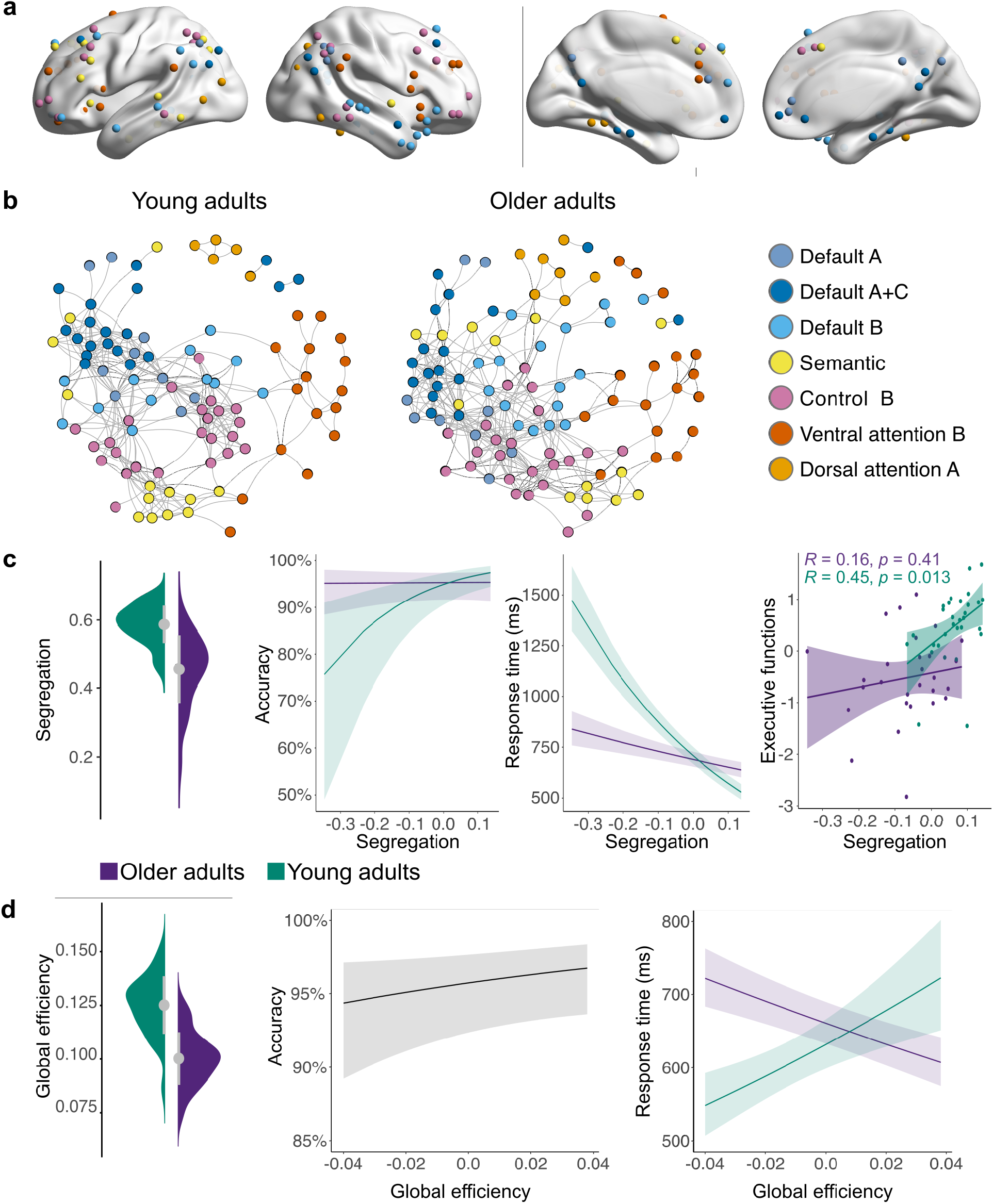
Age-related differences in whole-brain segregation and integration and their behavioral relevance. (a) For each participant, a task-related brain network graph was constructed using 121 nodes. The nodes were based on significant global and local peak maxima of the seven networks derived from the ICA (see Table S1 for exact locations of nodes). (b) Spring-embedded graphs depicting age differences in the modular organization of the brain. Graphs are based on average connectivity in each age group. Stronger segregation is reflected by higher within- and lower between-network correlations. In comparison, young adults show stronger segregation than older adults for most networks. For visualization purposes, graphs are displayed at 5% graph density. (c) Brain-wide system segregation was higher for young adults and had distinct effects on behavior for each age group with young adults profiting from increasing segregation. (d) A different picture emerged for global efficiency. Global efficiency was calculated for individual orthogonal minimum spanning trees, which were based on weighted correlation matrices. The graphs of young adults showed stronger global efficiency than older adults. While increasing global efficiency was associated with better performance in both age groups, it predicted slower performance in young and faster performance in older adults. Note that segregation and global efficiency values were mean-centered for analyses with behavior.

We used the measure of global efficiency to assess network integration. A linear mixed-effects model indicated higher global efficiency for young adults (*x*^2^= 21.86, *p* < 0.001; Figure 5d; Table S9). Efficiency values were then entered into regression models to assess their predictive value. Results showed a significant main effect of global efficiency for accuracy (*x*^2^= 8.79, *p* = 0.003) and a significant interaction with age for response time (*x*^2^= 41.79, *p* < 0.001). While increasing system-wide efficiency was generally associated with better performance, it also predicted faster performance in older adults but slower responses in young adults (Figure 5d; Table S10).

### Brain system segregation predicts age-related differences in behavior as a function of network type

We next examined whether segregation differed between networks. Previous research showed that networks exhibit differences in their patterns of age-related changes in segregation (Chan et al., 2014). While these studies focused on a broad distinction of sensorimotor and cognitive association networks, we investigated segregation and its behavioral relevance for each network individually to explore age-accompanied differences as a function of system type. Overall, results showed that all networks were less segregated in older than young adults (*x*^2^= 47.06, *p* < 0.001; Figure 6a; Table S11). However, networks’ increasing segregation differed in their behavioral relevance (Table S12).

**Figure 6.**
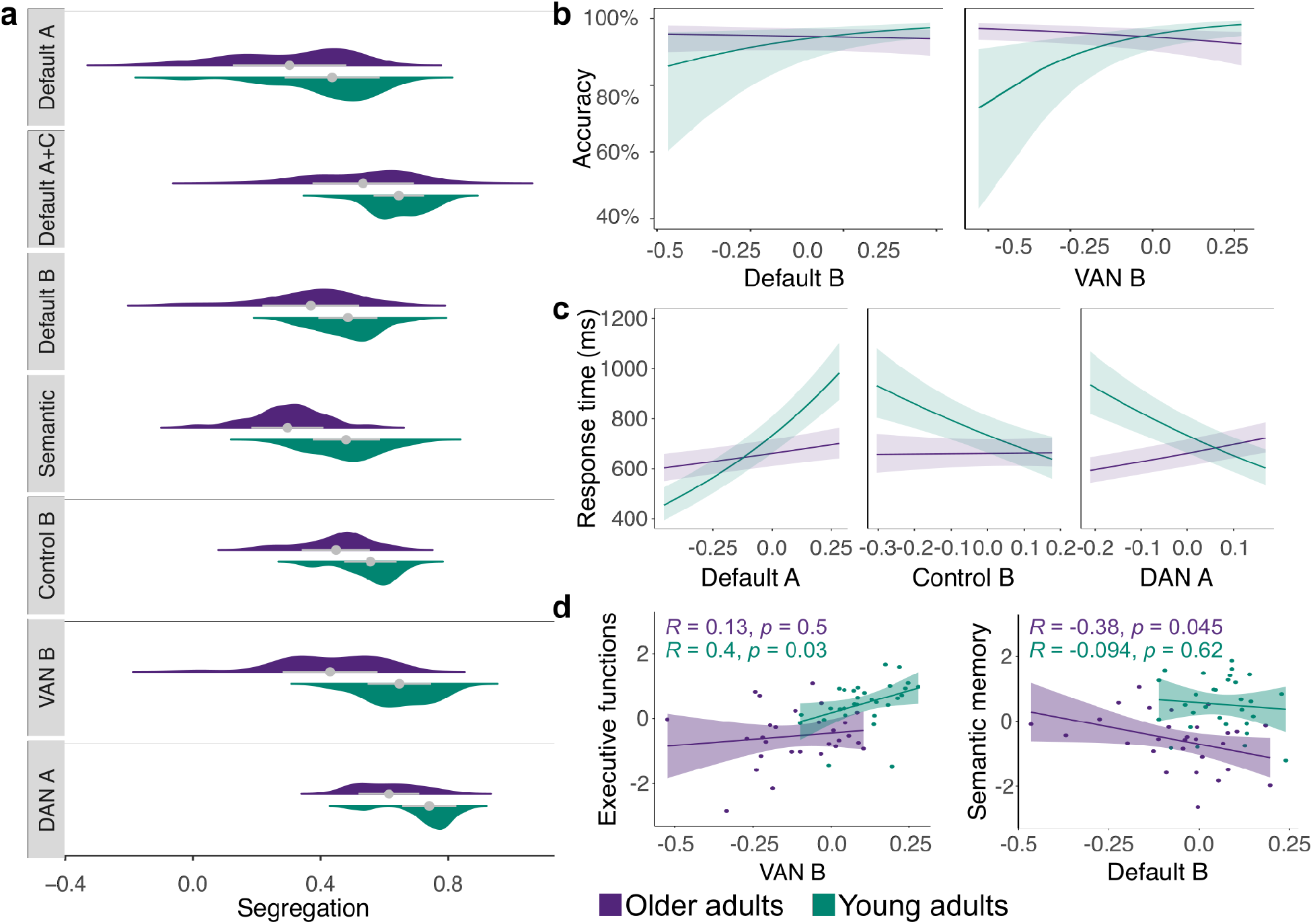
Segregation of individual networks is associated with distinct behavior of older and young adults as a function of system type. (a) Individual networks’ segregation values by age. All networks showed stronger segregation in young adults. (b) Generalized linear mixed-effects models for accuracy revealed significant interactions with age and network segregation for two systems while (c) linear mixed-effects models for response time showed significant interactions for three networks. For most networks, increasing segregation was associated with better and faster performance in young adults and worse and slower reactions in older adults. (d) Significant correlations between network segregation and neuropsychological measures. For young adults, we detected a positive correlation of increasing segregation of VAN-B with executive functions, whereas for older adults, a negative correlation of increasing segregation of DMN-B with semantic memory was found. Note that segregation and global efficiency values were mean-centered for analyses with behavior. VAN ventral attention network, DAN dorsal attention network.

For accuracy, we detected significant interactions between age and network segregation (Figure 6b) for DMN-B (*x*^2^= 5.76, *p* = 0.016) and VAN-B (*x*^2^= 18.22, *p* < 0.001). For response time (Figure 6c), results showed significant interactions with age and the networks DMN-A (*x*^2^= 79.3, *p* < 0.001), CONT-B (*x*^2^= 21.16, *p* < 0.001), and DAN-A (*x*^2^= 68.62, *p* < 0.001). Overall, stronger segregation of different systems was associated with better and faster performance for young adults and poorer and slower reactions in older adults. Only increasing segregation of DMN-A predicted slower reactions in young adults which might point to a different role of this system in semantic cognition.

We also explored the relationship of network segregation with neuropsychological measures (Figure 6d). Results revealed a significant positive correlation of segregation in the VAN-B with executive measures in young adults (*r* = 0.4, *p* = 0.03) and a negative correlation of DMN-B with semantic memory in older adults (*r* = −0.38, *p* = 0.045).

In summary, exploring brain system integration and segregation in a semantic task revealed age-specific dynamics where young adults clearly profit from a stronger modular network organization whereas increasing integration improves efficiency only in the aging brain.

### Stronger system-wide integration of brain networks in older adults is facilitated by additional connector hubs in frontal and temporal regions

An important characteristic of large-scale brain organization is the presence of regions, or nodes, that play an important role in facilitating communication between communities of a network. These nodes, commonly referred to as connector hubs, are defined by a high number of connections (edges) diversely distributed across communities (Bertolero et al., 2017). Previous work has highlighted their crucial role for integrative processing in resting- and task-state networks (Cohen and D’Esposito, 2016). We explored the existence of connector hubs via the normalized participation coefficient (PC (Pedersen et al., 2020). Results revealed connector hubs in bilateral frontal, parietal, and temporal regions in both age groups (Figure 7a; Table S13-14). Notably, there were multiple nodes from the subsystems of the default network and CONT-B identified as connector hubs in the bilateral regions of the inferior parietal lobe and AG. Furthermore, both age groups had connector hubs in the right MTG and MFG. In older adults, additional connector hubs were found in the left inferior temporal gyrus and the frontal pole. A linear model revealed nodes with stronger PC only in the graphs of older adults: in the frontal pole, which was also identified as a connector hub, STG, and bilateral fusiform gyri (Figure 7b; Table S15).

**Figure 7.**
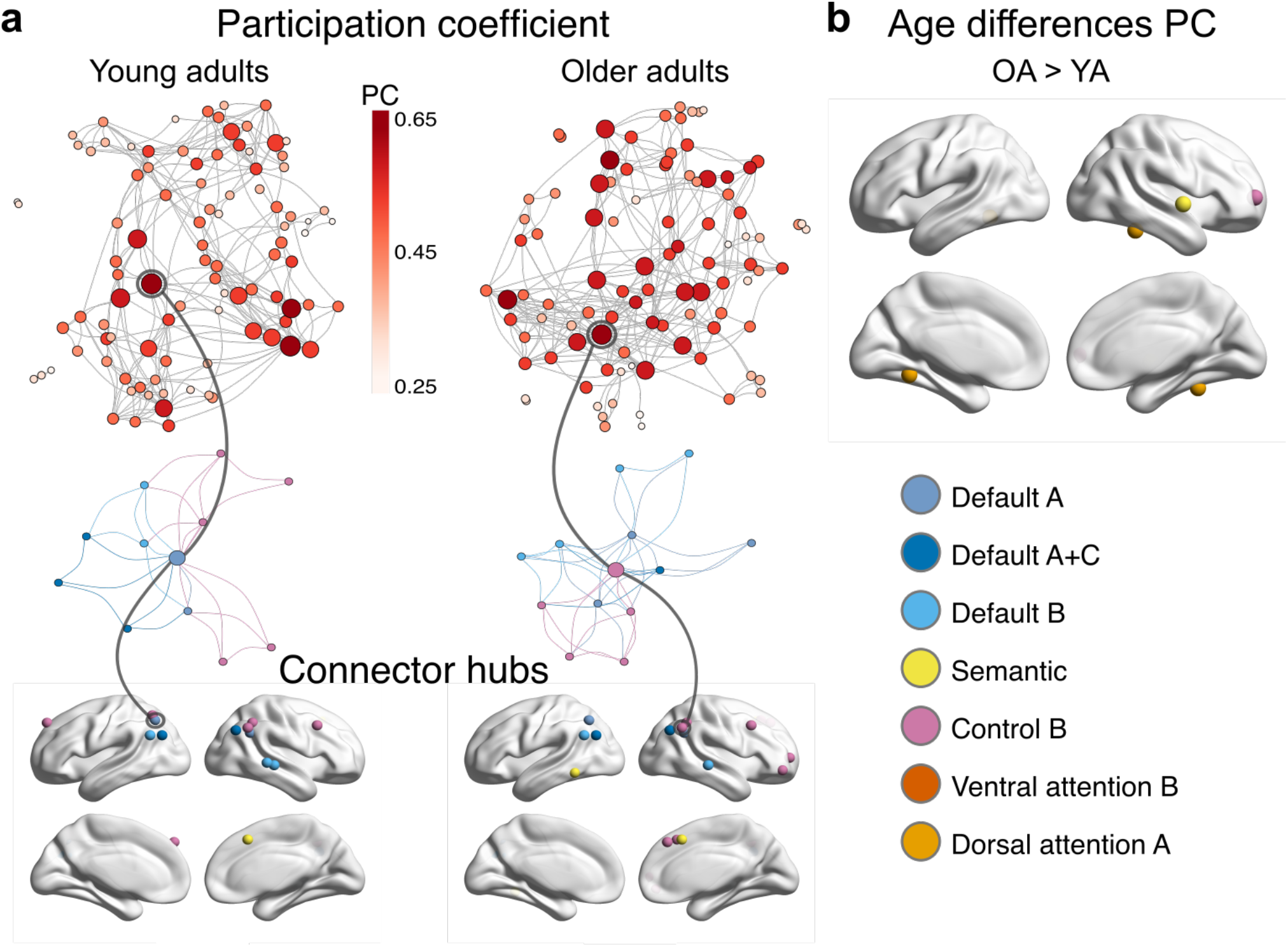
Topology of network hubs in young and older adults. (a) The normalized participation coefficient (PC) was calculated for individual orthogonal minimum spanning trees (OMST). Graphs display the PC of each node for the average OMST in each age group (top). For visualization purposes, the strongest 5% of connections are shown. Stronger PC values are reflected by color and node size. The higher the PC, the more a node is connected with nodes from other communities. The node with the highest PC value in each age group is extracted and displayed with its neighboring nodes colored by community (middle). Note that these connector hubs are connected to many different communities. Connector hubs were defined in each age group via PC values at least 1SD above the mean. In both groups, connector hubs were detected in frontal, parietal, and temporal regions (bottom). (b) A linear model with age as predictor revealed nodes with stronger PC only in older adults. The top and middle graphs were plotted using the ForceAtlas2 algorithm. The force-directed layout causes nodes of the same community to cluster together and diversely connected hubs (connector hubs) to appear in the center of the graph.

## Discussion

The neural bases of cognitive aging remain poorly understood. It is especially debated how age-related neural reorganization impacts cognition. A better understanding of the neural resources that help to maintain cognitive functions and counteract decline would be mandatory to design efficient treatment and training protocols. In the present study, we approached this unresolved issue by investigating the functional connectome of young and older adults in semantic cognition, a key domain of human cognition largely preserved in healthy aging. Our results demonstrate a reconfigured network architecture with age even when word retrieval abilities remain intact. Overall, networks showed increased integration of task-negative and task-positive networks with age which manifested as increased coupling between functional connectivity networks, reduced segregation of global brain systems, and a larger number of connector hubs in brain graphs of older adults. Associating these network profiles with behavior revealed intact, albeit less efficient, performance for more integrated systems in older adults. These findings shed new light on the frequently reported pattern of declining brain system segregation with age and its impact on cognition (Chan et al., 2014; Geerligs et al., 2014). Our results indicate a compensatory role of increased brain system integration but also reveals its limitations in terms of economical processing.

Using task-based fMRI data and group spatial ICA, we characterized seven higher-order large-scale functional networks relevant to semantic word retrieval. These included default, semantic, frontoparietal control, and attention networks. Notably, our analysis detected two networks associated with semantic processing: a network component that showed strong activity in frontal regions that have been attributed to semantic control processes (Jackson, 2021) and another component overlapping with the subnetwork DMN-B (Yeo et al., 2011), which has been proposed to facilitate access to semantic knowledge (Smallwood et al., 2021). Thus, these two semantic subnetworks appear to represent complementary aspects of semantic cognition. Moreover, in line with our previous work (Martin et al., 2022), we detected default and cognitive control systems, lending support to the notion that networks that have been characterized as anti-correlated during restingstate become functionally integrated for successful task processing when controlled access to semantic memory is required (Krieger-Redwood et al., 2016). Indeed, exploring task-based functional connectivity showed strong positive coupling between distinct cognitive networks in both age groups. Two subnetworks of the default network, DMN-A and DMN-B, were strongly coupled with the frontoparietal control network. This finding agrees with accumulating evidence that the default network integrates with control and executive resources during goal-directed task processing (Krieger-Redwood et al., 2016), especially when complex behavior is supported by knowledge (Wang et al., 2021), and thus enables flexible cognition (Smallwood et al., 2021).

Examining age-related differences in network coupling revealed additional integration of distinct networks with age. Older adults showed stronger positive coupling of SEM, CONT-B, and DAN-A with VAN-B relative to young adults, suggesting an increased cognitive demand during semantic processing. In contrast, networks of young adults displayed stronger decoupling of default with attention networks. Previous work indicates that young adults can benefit from a more integrated brain organization in situations of high task demand to facilitate information flow across components (Vatansever et al., 2015; Cohen and D’Esposito, 2016; Zhang et al., 2020). Our results transfer this observation to the aging brain and demonstrate increased crosstalk between networks with age. Importantly, when we associated the differences in network coupling with behavior, we found that enhanced coupling of different cognitive systems like default and attention networks was associated with consistently high but less efficient performance in older adults. Consistent with results from different cognitive domains (Gallen et al., 2016; Adnan et al., 2019; Crowell et al., 2020; Deng et al., 2021), we demonstrate that enhanced network integration with age is linked to high accuracy but at the cost of efficiency. While such reorganization helps older adults to maintain cognitive flexibility, it might not be the most efficient form of wiring.

We applied graph theory to further explore age-accompanied changes in the network architecture. Our results revealed global decreases in segregation and efficiency with age. The reduction of segregation in older adults is in line with previous work from resting-state (Chan et al., 2014; Sala-Llonch et al., 2014; Geerligs et al., 2015) and task-based studies (Geerligs et al., 2014; Gallen et al., 2016; Crowell et al., 2020), as well as longitudinal investigations (Betzel et al., 2014; Cao et al., 2014; Chong et al., 2019), and suggests that aging is associated with a reduced ability for specialized processing within highly connected clusters (Rubinov and Sporns, 2010). This was confirmed by our results on segregation of each individual network where young adults generally showed stronger segregation.

In terms of global efficiency, most studies reported lower global efficiency in older adults (Sala-Llonch et al., 2014; Chong et al., 2019; Gonzalez-Burgos et al., 2021), although others have also reported no changes or even increases with age (Cao et al., 2014; Chan et al., 2014; Song et al., 2014; Geerligs et al., 2015). These discrepancies might stem from methodological considerations such as the number of nodes in a brain graph since global efficiency is based on the length of its edges (Zalesky et al., 2010) or different thresholding methods of connectivity matrices like the commonly applied proportional thresholding, which has been shown to introduce spurious correlations and inflate group-related differences in graph metrics (van den Heuvel et al., 2017). To avoid these pitfalls, our calculation of global efficiency was based on the recently developed OMST (Dimitriadis et al., 2017), a data-driven approach of individualized graph construction with high reliability (Jiang et al., 2021; Luppi and Stamatakis, 2021).

Reduced global efficiency implies higher wiring cost and a less efficient information flow among distributed networks of the global brain system (Bullmore and Sporns, 2012). This is especially relevant for the processing of complex cognitive functions like semantic word retrieval which require the integration of distinct networks, as revealed by our functional connectivity analyses. At the neurobiological level, these changes have been associated with reduced functional connectivity of long-range connections in older adults (Sala-Llonch et al., 2014). Thus, even though functional networks become more integrated with age, potentially due to stronger activation of more but less specialized nodes, the efficient information transfer between networks is impaired leading to slower processing in aging. This observation may represent an overall decline of cognitive attention systems in the aging brain, reflected in slower responses with similar task accuracy, which was already evident at the behavioral level in our data.

Additional evidence for this interpretation stems from the larger number of connector hubs in older adults. In the young brain, an increase of connector hubs has been linked to enhanced task demands to facilitate integration across different networks and enable better task performance (Bertolero et al., 2018; Zhang et al., 2020). Resting-state studies in healthy aging also reported more connector hubs, indicating a reduced distinctiveness of network-specific nodes (Geerligs et al., 2015; Chan et al., 2017; Chong et al., 2019). Our work confirms these findings during task processing and allows an interpretation in light of the semantic nature of our task. Nodes with a higher participation coefficient in older adults were located in frontal and temporal regions and associated with CONT-B, DAN-A, and SEM networks. This result underlines the enhanced cognitive demand during semantic word retrieval with age and provides a mechanistic explanation for the frequently reported pattern of over-activation of prefrontal control regions during demanding task processing in older adults (Davis et al., 2008). A reduced selectivity in activation of network nodes and hence an over-recruitment of less specialized brain regions leads to a decline in efficient neural processing between brain regions, and this process might form the basis of neural dedifferentiation in aging (Chan et al., 2017; Chong et al., 2019). Its effect on cognition, aberrant or compensatory, depends on the neurocognitive requirements of a task.

Exploring the topology of task-relevant neural networks as a function of cognitive performance allowed us to directly link observed age-related differences with behavior. Results showed that young adults strongly capitalized on a more segregated system during task processing in the form of faster and better performance. In contrast, increasing whole-brain segregation predicted faster but not better performance in older adults, whereas increasing global efficiency predicted better performance across groups but faster responses only in older adults. These findings have important implications for current theories on the behavioral impact of network reorganization in aging. While a less selective and more integrated network organization might not be the most efficient system in terms of processing speed, it enables older adults to maintain high performance. However, stronger integration does not automatically imply a more efficient system as evident by a generally reduced global efficiency in brain graphs of older adults and a predicted faster response in a more efficient system. Our findings lend support to a compensatory mechanism of age-accompanied reconfiguration in network topologies. Importantly, they also reveal the limitations of such compensatory reorganization processes and demonstrate that a youth-like network architecture in terms of balanced integration and segregation is associated with more economical processing.

## Acknowledgments

SM held a stipend by the German Academic Scholarship Foundation (Studienstiftung des deutschen Volkes). DS was supported by the Deutsche Forschungsgemeinschaft (SA 1723/5-1) and the James S. McDonnell Foundation (Understanding Human Cognition, #220020292). GH was supported by the Lise Meitner excellence program of the Max Planck Society and the Deutsche Forschungsgemeinschaft (HA 6314/3-1, HA 6314/4-1). The authors would like to thank the medical technical assistants of MPI CBS for their support with data acquisition, and Annika Dunau, Caroline Duchow, and Rebekka Luckner for their support with transcriptions of recordings.

