## Supplementary Materials for "Age-related reorganization of functional network architecture in semantic cognition"

### Age-related reorganization of functional network architecture for language processing

### Supplementary Figures

#### Figure S1

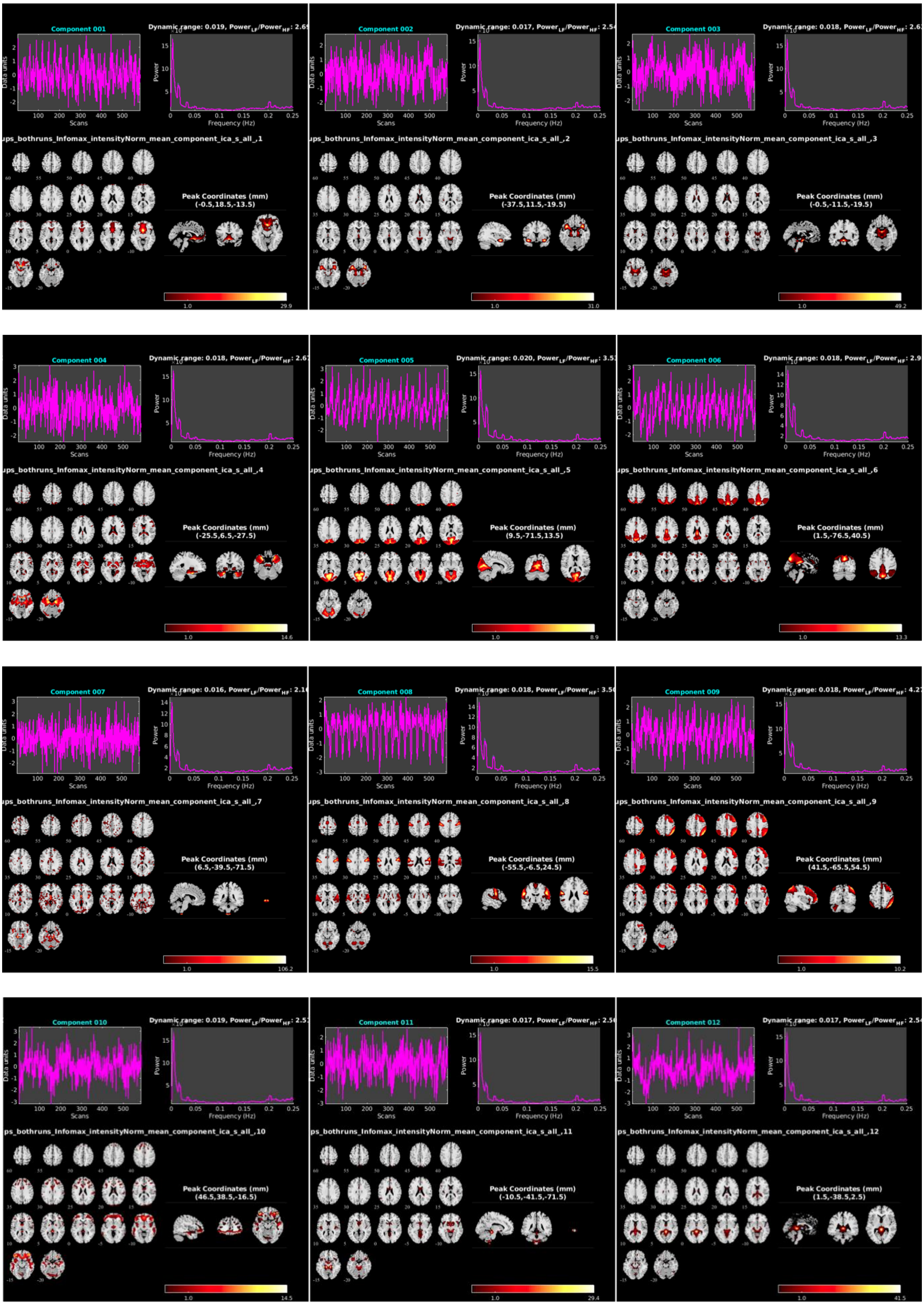

Supplementary Figures

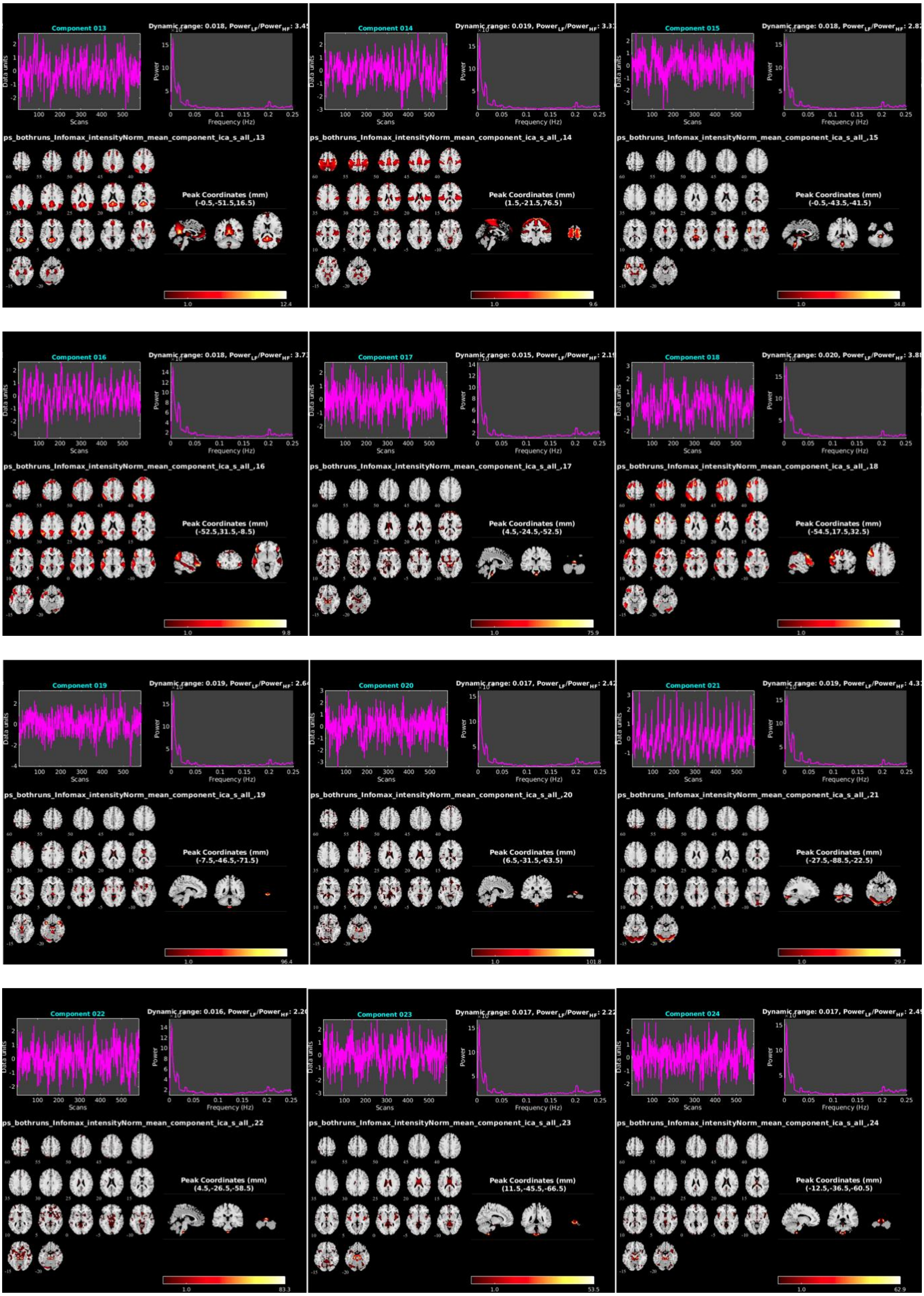

Supplementary Figures

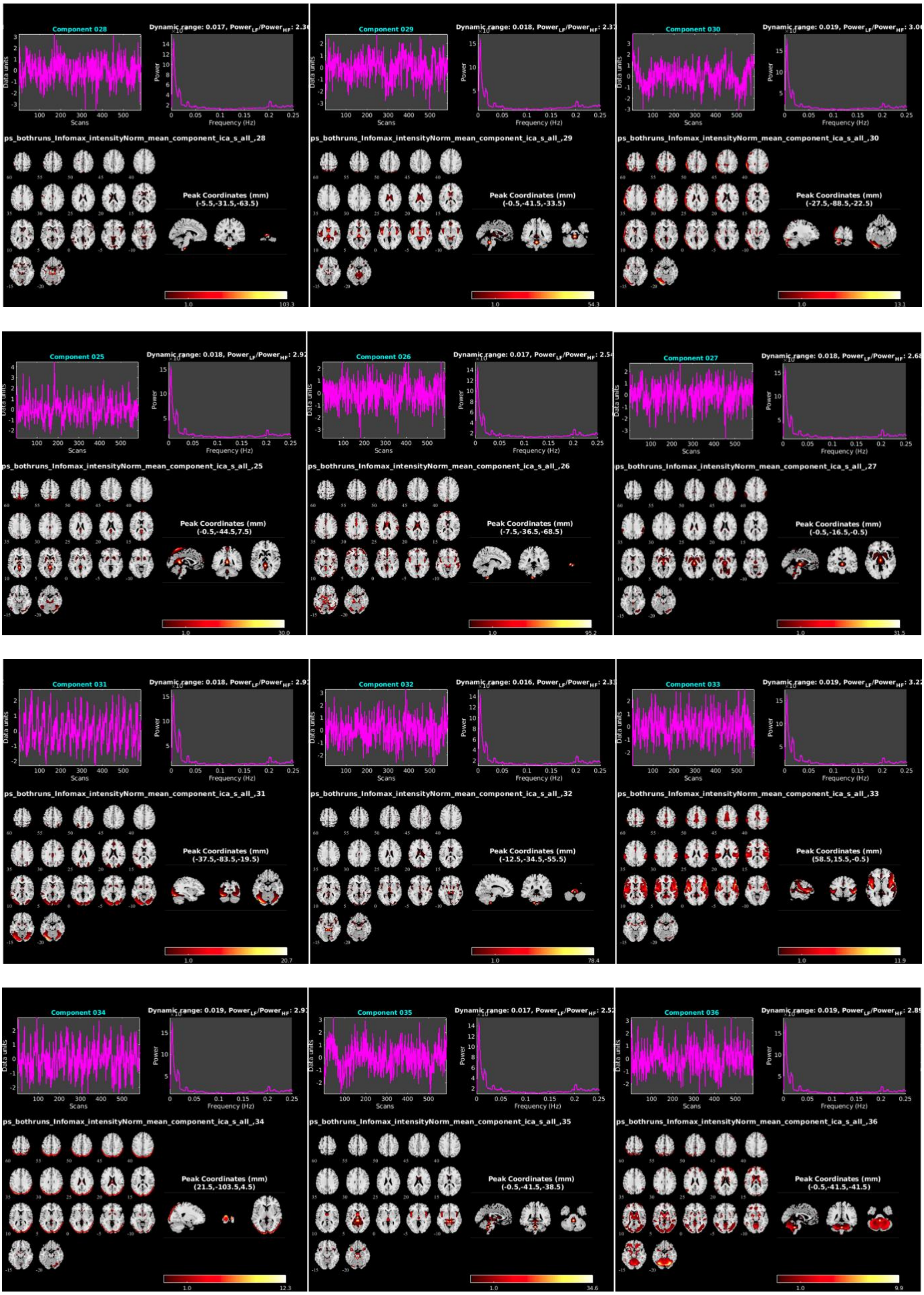

Supplementary Figures

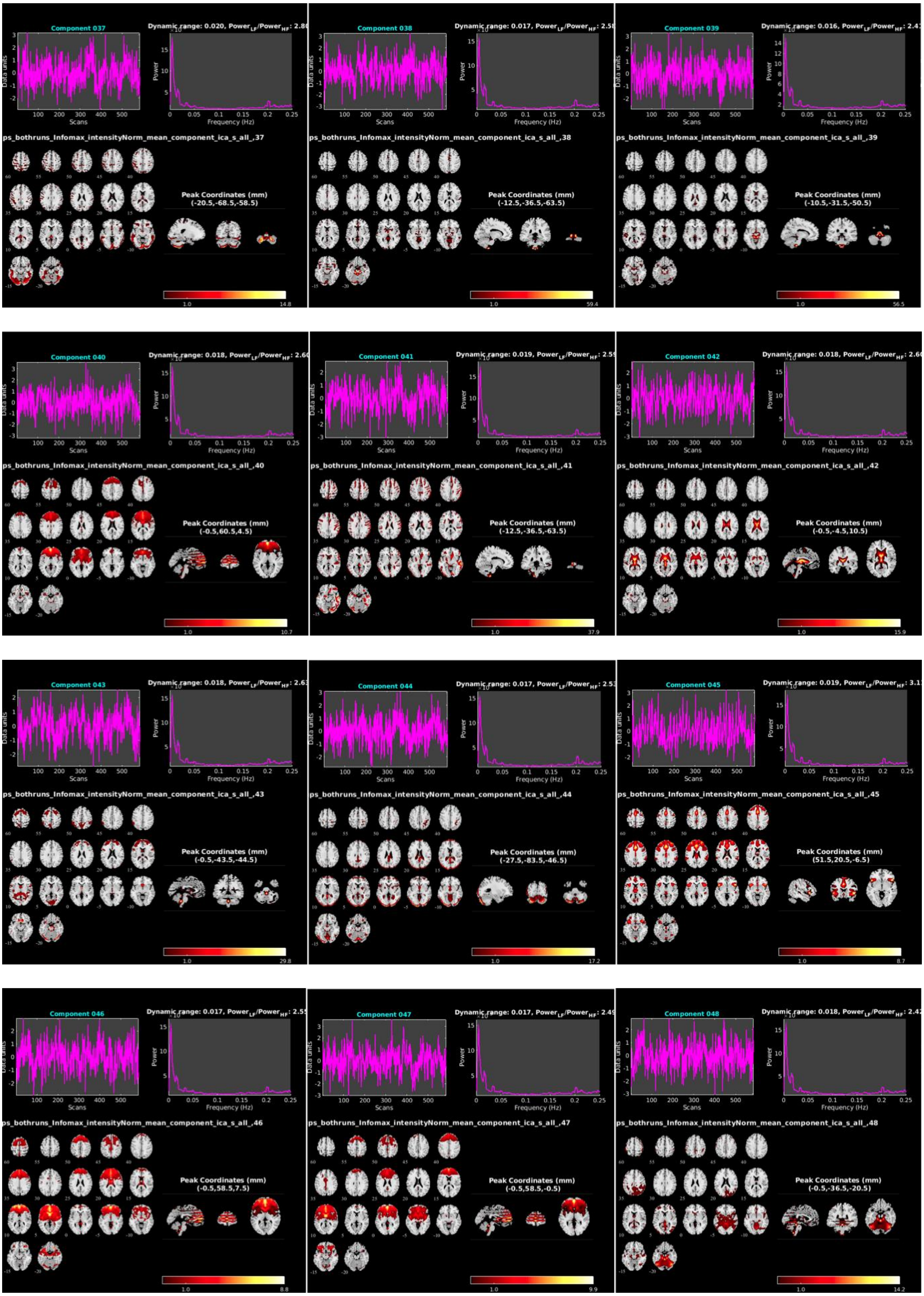

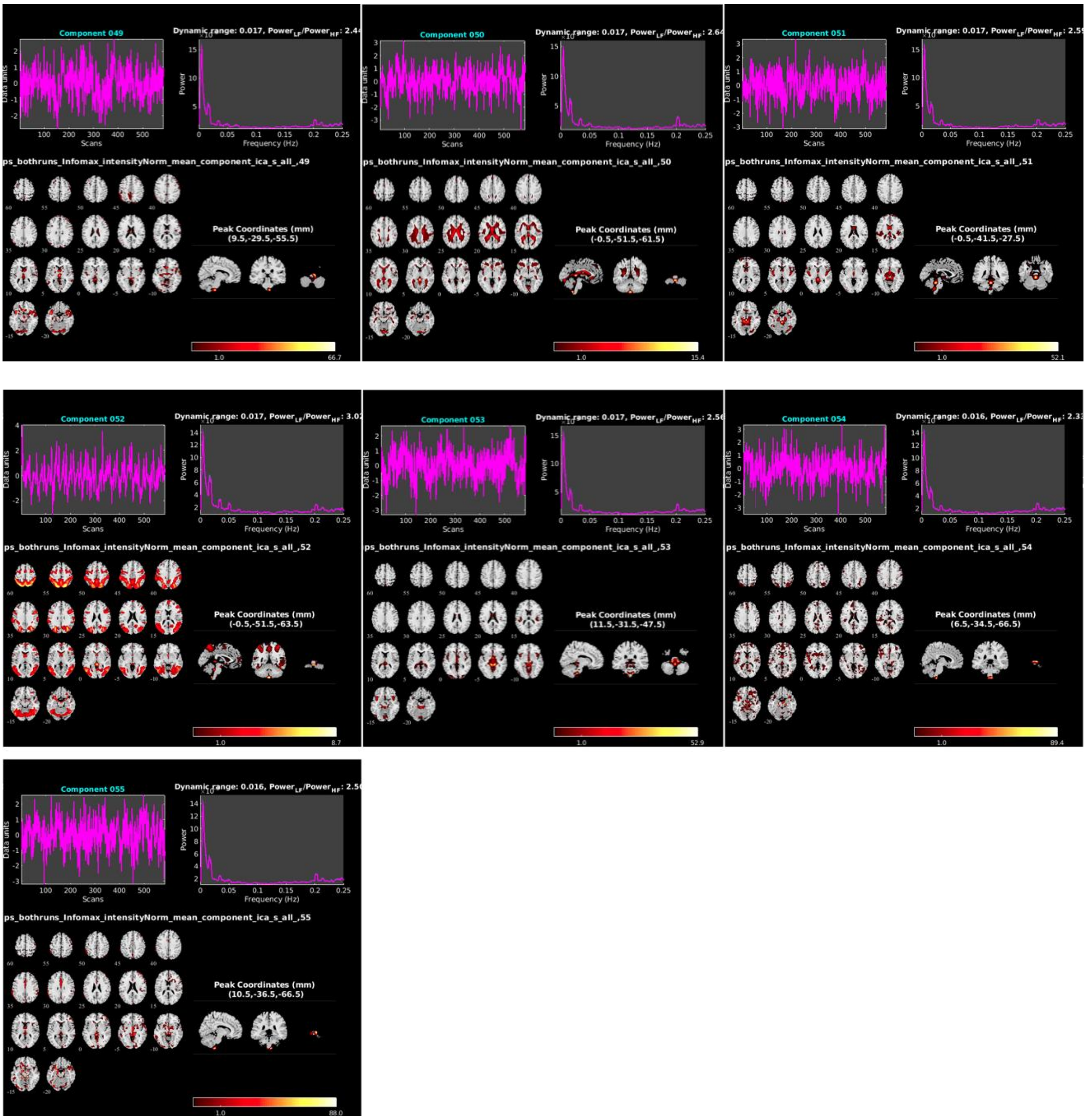

**Figure S1. 55 independent network components derived via group spatial independent component analysis (ICA).** ICA was conducted using the Group ICA of fMRI Toolbox. Dimensions were reduced to 55 using minimum description length information criteria. Icasso was repeated 50 times to ensure reliability of the decomposition, and group-level ICs were back-reconstructed to the participant level using the group-information guided ICA (GICA3) algorithm.

**Figure S2**

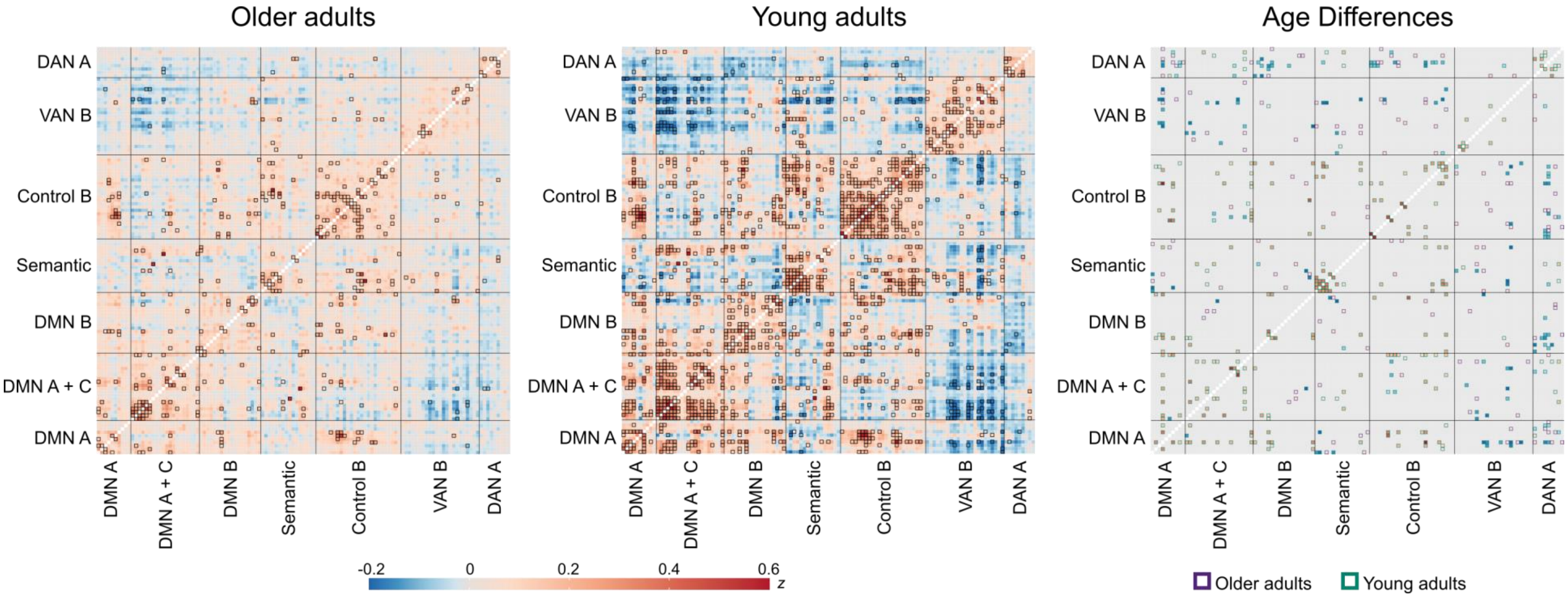

**Figure S2. Heatmaps display significant results of functional coupling between the regions of interest (n = 121) of the ICA-derived networks.** Connectivity values are Fisher-transformed partial correlations. Bold frames indicate significant values which are based on cPPI-derived significance values in the age groups while age differences were assessed using permutation testing in network-based statistics (cluster-forming threshold at p = 0.01, FWE-corrected significance threshold at p = 0.05 with 10,000 permutations).

**Figure S3**

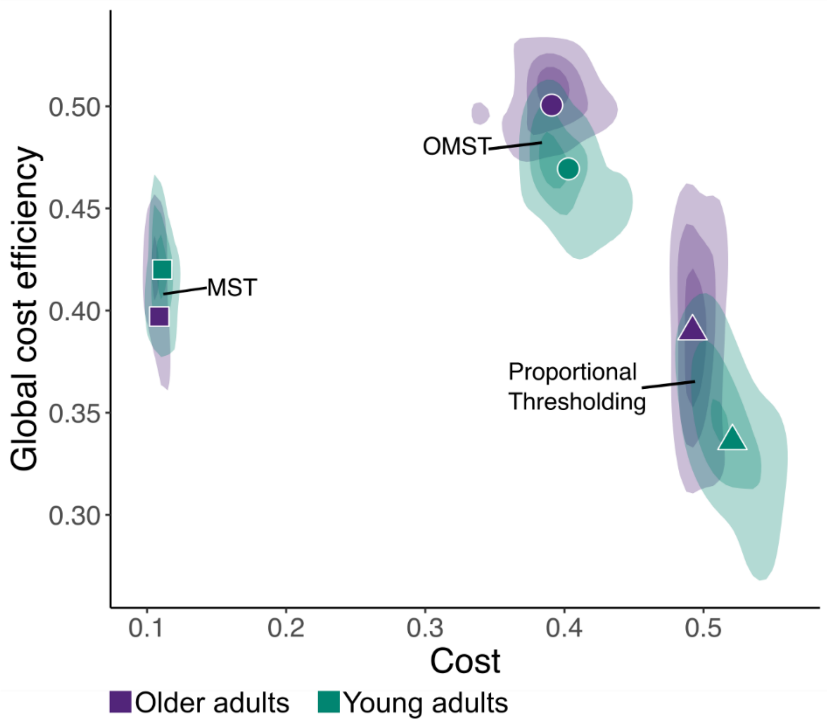

**Figure S3. Plots of global cost efficiency (GCE) against cost for different filtering options of graphs.** We compared the GCE of two topological (minimum spanning tree (MST) and orthogonal minimum spanning tree (OMST)) and one proportional (5-20% strongest edge weights) edge filtering method in both age groups. Symbols indicate the average value in each group while densities display value distribution.

**Table S1.** Characteristics and neuropsychological test results of participants

| | Young adults ($n$ = 30) | Older adults ($n$ = 28) |
|---|---|---|
| **Demographics** | | |
| Age (years) | 27.6 (4.4) | 65.2 (2.8) |
| Gender (F:M) | 16:14 | 14:14 |
| Education (years) | 18.7 (2.6) | 15.2 (2.5)* |
| Beck Depression Inventory (cut-off 18 points) | – | 4.7 (4.1) |
| **Neuropsychological** | | |
| Spot-the-word test (max. 40) | 29.1 (3.2) | 31.5 (2.5)* |
| Semantic fluency (sum surnames, hobbies) | 51.2 (8.4) | 40.7 (6.7)* |
| Reading span test (max. 6) | 3.5 (1) | 2.9 (0.7)* |
| Digit symbol substitution test (max. 90 in 90 s) | 72.1 (11.4) | 50.2 (10.4)* |
| Trail Making Test A (time in s) | 17.3 (5.8) | 25.4 (6.4)* |
| Trail Making Test B (time in s) | 36.1 (11.9) | 61.8 (29.4)* |
| Mini-Mental State Examination (max. 30 points) | – | 28.36 (1.2) |

*Note.* Mean values of raw scores with standard deviations. * Significant differences between age groups at $p < 0.01$.

**Table S2.** Behavioral results of mixed-effects models for accuracy and response time

| | Accuracy | | | Response time | | |
|---|---|---|---|---|---|---|
| *Coefficient* | *Log-Odds* | *CI* | *p* | *Estimates* | *CI* | *p* |
| Intercept | 4.61 | 3.68 – 5.55 | **<0.001** | 6.43 | 6.39 – 6.48 | **<0.001** |
| Age | -0.50 | -0.93 – -0.08 | **0.021** | 0.01 | 0.00 – 0.03 | **0.043** |
| Condition | -3.06 | -4.89 – -1.22 | **0.001** | 0.15 | 0.08 – 0.22 | **<0.001** |
| Education | -0.11 | -0.23 – 0.01 | 0.082 | -0.01 | -0.01 – 0.00 | 0.059 |
| Age * Condition | 0.75 | -0.06 – 1.55 | 0.069 | 0.08 | 0.06 – 0.10 | **<0.001** |
| **Random Effects** | | | | | | |
| $\sigma^2$ | 3.29 | | | 0.09 | | |
| $\tau_{00}$ | 0.19 $_{Subj}$ | | | 0.01 $_{Subj}$ | | |
| | 1.52 $_{Category}$ | | | 0.00 $_{Category}$ | | |
| ICC | 0.34 | | | 0.10 | | |
| Observations | 19710 | | | 18877 | | |
| Marginal $R^2$ / Conditional $R^2$ | 0.328 / 0.558 | | | 0.055 / 0.149 | | |

*Note*. Significant effects are marked in bold. Contrasts are sum coded. P-values were obtained via likelihood ratio tests. CI Confidence interval.

**Table S3.** Jaccard similarity coefficients with the 7-networks parcellation scheme

| | **IC06** | **IC09** | **IC13** | **IC16** | **IC18** | **IC45** | **IC52** |
|---|---|---|---|---|---|---|---|
| Control | 0.128 | **0.287** | 0.039 | 0.054 | *0.194* | *0.161* | 0.090 |
| Default | **0.155** | *0.154* | **0.234** | **0.412** | 0.133 | 0.133 | 0.043 |
| Dorsal attention | 0.073 | 0.089 | 0.067 | 0.022 | 0.139 | 0.059 | **0.301** |
| Limbic | 0.000 | 0.004 | 0.012 | 0.014 | 0.008 | 0.005 | 0.011 |
| Ventral attention | 0.039 | 0.045 | 0.025 | 0.039 | 0.053 | **0.211** | 0.097 |
| Somatomotor | 0.017 | 0.064 | 0.029 | 0.057 | 0.067 | 0.044 | 0.052 |
| Visual | 0.038 | 0.015 | 0.090 | 0.046 | 0.045 | 0.028 | *0.177* |
| General semantic cognition | 0.032 | 0.030 | 0.072 | *0.194* | **0.201** | 0.092 | 0.050 |
| Semantic control | 0.012 | 0.036 | 0.012 | 0.067 | *0.153* | 0.091 | 0.027 |

*Note.* The selected network labels for the respective independent components are shown in bold while all cognitive networks that showed a higher similarity coefficient than $J$ = 0.15 are shown in italics.

**Table S4.** Significant clusters of task-relevant independent components

| Anatomical structure | Hemi | *k* | *t* | *x* | *y* | *z* |
|---|---|---|---|---|---|---|
| **IC 06** | | | | | | |
| **Precuneus Cortex** | **R** | **1057** | **19.94** | **5** | **-71** | **35** |
| Precuneus Cortex | L | | 17.67 | -5 | -66 | 38 |
| Cingulate Gyrus, posterior division | R | | 17.43 | 2 | -41 | 35 |
| **Angular Gyrus** | **L** | **242** | **14.56** | **-45** | **-64** | **46** |
| Angular Gyrus | L | | 13.29 | -48 | -56 | 44 |
| **Angular Gyrus** | **R** | **289** | **14.31** | **47** | **-56** | **44** |
| Angular Gyrus | R | | 10.44 | 54 | -51 | 41 |
| Angular Gyrus | R | | 9.58 | 52 | -54 | 33 |
| **Cingulate Gyrus, anterior division** | **R** | **51** | **7.58** | **2** | **43** | **5** |
| Paracingulate Gyrus | R | | 6.17 | 5 | 46 | -3 |
| **Paracingulate Gyrus** | **L** | **25** | **6.96** | **-3** | **38** | **22** |
| **Frontal Pole** | **L** | **16** | **6.43** | **-23** | **66** | **5** |
| **IC 09** | | | | | | |
| **Frontal Pole** | **R** | **564** | **21.83** | **42** | **48** | **-9** |
| Frontal Pole | R | | 20.52 | 49 | 43 | -9 |
| Frontal Pole | R | | 19.22 | 29 | 56 | 5 |
| **Middle Frontal Gyrus** | **R** | **810** | **21.33** | **49** | **31** | **33** |
| Middle Frontal Gyrus | R | | 18.52 | 37 | 21 | 55 |
| Middle Frontal Gyrus | R | | 18.25 | 52 | 16 | 41 |
| **Angular Gyrus** | **R** | **520** | **20.75** | **52** | **-51** | **44** |
| Angular Gyrus | R | | 20.72 | 57 | -56 | 38 |
| Angular Gyrus | R | | 20.26 | 39 | -61 | 52 |
| **Middle Temporal Gyrus, posterior division** | **R** | **104** | **18.62** | **62** | **-36** | **-12** |
| Middle Temporal Gyrus, temporooccipital part | R | | 16.94 | 59 | -46 | -12 |
| Middle Temporal Gyrus, temporooccipital part | R | | 14.67 | 67 | -39 | -3 |
| **Superior Frontal Gyrus** | **R** | **335** | **18.38** | **5** | **36** | **44** |
| Paracingulate Gyrus | R | | 18.04 | 7 | 26 | 46 |
| Superior Frontal Gyrus | L | | 10.93 | -5 | 33 | 46 |
| **Angular Gyrus** | **L** | **364** | **15.31** | **-48** | **-54** | **55** |
| Angular Gyrus | L | | 12.43 | -45 | -61 | 52 |
| Angular Gyrus | L | | 11.15 | -38 | -59 | 46 |
| **Frontal Pole** | **L** | **231** | **14.18** | **-43** | **53** | **2** |
| Frontal Pole | L | | 13.29 | -38 | 61 | -1 |
| Frontal Pole | L | | 10.39 | -48 | 46 | -12 |
| **Middle Frontal Gyrus** | **L** | **200** | **10.80** | **-50** | **26** | **35** |
| Middle Frontal Gyrus | L | | 10.27 | -48 | 21 | 44 |
| **Middle Temporal Gyrus, posterior division** | **L** | **41** | **7.73** | **-63** | **-36** | **-9** |
| **Superior Frontal Gyrus** | **L** | **10** | **6.08** | **-23** | **18** | **57** |
| **IC 13** | | | | | | |
| **Precuneus Cortex** | **L** | **1459** | **32.20** | **-5** | **-59** | **19** |

| Anatomical structure | Hemi | *k* | *t* | *x* | *y* | *z* |
|---|---|---|---|---|---|---|
| Precuneus Cortex | R | | 32.11 | 5 | -59 | 27 |
| Precuneus Cortex | R | | 26.92 | 7 | -54 | 16 |
| **Angular Gyrus** | **L** | **525** | **20.64** | **-48** | **-71** | **30** |
| Angular Gyrus | L | | 18.65 | -40 | -66 | 27 |
| Angular Gyrus | L | | 14.87 | -38 | -79 | 41 |
| **Parahippocampal Gyrus, posterior division** | **L** | **207** | **18.74** | **-25** | **-31** | **-17** |
| Parahippocampal Gyrus, anterior division | L | | 10.87 | -20 | -22 | -20 |
| Lingual Gyrus | L | | 10.86 | -25 | -41 | -6 |
| **Superior Frontal Gyrus** | **L** | **228** | **16.63** | **-23** | **28** | **49** |
| **Angular Gyrus** | **R** | **461** | **15.89** | **52** | **-64** | **24** |
| Angular Gyrus | R | | 13.36 | 54 | -69 | 35 |
| Angular Gyrus | R | | 11.74 | 44 | -54 | 22 |
| **Frontal Medial Cortex** | **L** | **718** | **15.37** | **-5** | **51** | **-9** |
| Cingulate Gyrus, anterior division | R | | 13.77 | 2 | 33 | -9 |
| Frontal Medial Cortex | R | | 12.56 | 2 | 53 | -6 |
| **Middle Temporal Gyrus, anterior division** | **R** | **109** | **14.58** | **64** | **1** | **-17** |
| **Parahippocampal Gyrus, posterior division** | **R** | **133** | **11.99** | **27** | **-31** | **-17** |
| Parahippocampal Gyrus, anterior division | R | | 11.68 | 22 | **-19** | -23 |
| **Superior Frontal Gyrus** | **R** | **185** | **11.08** | **22** | **31** | **46** |
| **IC 16** | | | | | | |
| **Frontal Pole** | **L** | **1390** | **23.11** | **-5** | **48** | **46** |
| Superior Frontal Gyrus | R | | 21.03 | 14 | 31 | 57 |
| Paracingulate Gyrus | L | | 20.26 | -5 | 53 | 19 |
| **Middle Temporal Gyrus, anterior division** | **L** | **1382** | **23.08** | **-55** | **1** | **-20** |
| Middle Temporal Gyrus, posterior division | L | | 21.78 | -58 | -31 | -3 |
| Frontal Pole | L | | 19.89 | -50 | 43 | -12 |
| **Angular Gyrus** | **L** | **356** | **22.67** | **-55** | **-59** | **30** |
| Supramarginal Gyrus, posterior division | L | | 14.15 | -63 | -49 | 41 |
| **Inferior Frontal Gyrus, pars triangularis** | **R** | **313** | **17.26** | **52** | **31** | **-6** |
| Temporal Pole | R | | 12.61 | 37 | 23 | -23 |
| Frontal Orbital Cortex | R | | 11.27 | 44 | 23 | -14 |
| **Temporal Pole** | **R** | **238** | **16.84** | **52** | **16** | **-25** |
| Middle Temporal Gyrus, anterior division | R | | 16.64 | 52 | 3 | -31 |
| Temporal Pole | R | | 15.86 | 49 | 11 | -34 |
| **Middle Temporal Gyrus, posterior division** | **R** | **193** | **16.70** | **64** | **-29** | **-3** |
| Middle Temporal Gyrus, posterior division | R | | 15.08 | 54 | -24 | -6 |

| Anatomical structure | Hemi | *k* | *t* | *x* | *y* | *z* |
|---|---|---|---|---|---|---|
| Middle Temporal Gyrus, posterior division | R | | 14.76 | 67 | -36 | -1 |
| **Middle Frontal Gyrus** | **L** | **151** | **16.01** | **-40** | **16** | **52** |
| **IC 18** | | | | | | |
| **Insular Cortex** | **L** | **1554** | **28.83** | **-33** | **21** | **-1** |
| Inferior Frontal Gyrus, pars triangularis | L | | 21.00 | -48 | 28 | 19 |
| Middle Frontal Gyrus | L | | 19.65 | -50 | 21 | 33 |
| **Superior Frontal Gyrus** | **L** | **511** | **22.76** | **-5** | **31** | **44** |
| Paracingulate Gyrus | L | | 19.99 | -5 | 13 | 52 |
| Paracingulate Gyrus | R | | 18.29 | 2 | 21 | 46 |
| **Inferior Temporal Gyrus, temporooccipital part** | **L** | **984** | **18.99** | **-58** | **-49** | **-12** |
| Superior Temporal Gyrus, posterior division | L | | 10.54 | -60 | -31 | 5 |
| Parahippocampal Gyrus, posterior division | L | | 9.82 | -30 | -31 | -17 |
| **Angular Gyrus** | **L** | **270** | **16.99** | **-35** | **-59** | **41** |
| Angular Gyrus | L | | 16.88 | -30 | -66 | 49 |
| Angular Gyrus | L | | 16.52 | -30 | -71 | 41 |
| **Superior Frontal Gyrus** | **L** | **50** | **11.92** | **-23** | **26** | **46** |
| Superior Frontal Gyrus | L | | 9.63 | -15 | 36 | 46 |
| Frontal Pole | L | | 8.39 | -13 | 48 | 44 |
| **Temporal Fusiform Cortex, anterior division** | **L** | **22** | **9.50** | **-38** | **-9** | **-28** |
| **Superior Temporal Gyrus, anterior division** | **R** | **32** | **9.14** | **59** | **-4** | **-1** |
| **IC 45** | | | | | | |
| **Frontal Pole** | **L** | **583** | **23.44** | **-25** | **38** | **30** |
| Frontal Pole | L | | 21.05 | -23 | 46 | 24 |
| Frontal Pole | L | | 19.36 | -30 | 53 | 24 |
| **Paracingulate Gyrus** | **L** | **565** | **21.83** | **-5** | **31** | **33** |
| Cingulate Gyrus, anterior division | L | | 21.81 | -5 | 31 | 22 |
| Paracingulate Gyrus | R | | 21.76 | 14 | 28 | 27 |
| **Frontal Pole** | **R** | **630** | **20.80** | **27** | **48** | **30** |
| Frontal Pole | R | | 20.35 | 27 | 41 | 24 |
| Frontal Pole | R | | 18.56 | 37 | 43 | 30 |
| **Frontal Operculum Cortex** | **L** | **225** | **16.57** | **-35** | **16** | **11** |
| Frontal Orbital Cortex | L | | 15.46 | -33 | 26 | -6 |
| Inferior Frontal Gyrus, pars opercularis | L | | 15.04 | -50 | 13 | -3 |
| **Frontal Operculum Cortex** | **R** | **280** | **16.15** | **47** | **18** | **-3** |
| Frontal Operculum Cortex | R | | 15.26 | 39 | 18 | 11 |
| Frontal Orbital Cortex | R | | 14.28 | 39 | 18 | -12 |
| **Supramarginal Gyrus, posterior division** | **R** | **157** | **14.98** | **62** | **-44** | **27** |
| Supramarginal Gyrus, posterior division | R | | 13.20 | 67 | -39 | 35 |

| Anatomical structure | Hemi | *k* | *t* | *x* | *y* | *z* |
|---|---|---|---|---|---|---|
| Supramarginal Gyrus, posterior division | R | | 5.93 | 62 | -39 | 49 |
| **Superior Frontal Gyrus** | **L** | **27** | **14.57** | **-13** | **3** | **68** |
| Superior Frontal Gyrus | L | | 12.61 | -8 | 8 | 63 |
| **Supramarginal Gyrus, posterior division** | **L** | **41** | **11.77** | **-60** | **-44** | **27** |
| **Inferior Frontal Gyrus, pars opercularis** | **R** | **22** | **8.01** | **52** | **11** | **11** |
| Inferior Frontal Gyrus, pars opercularis | R | | 6.73 | 54 | 13 | 22 |
| **Frontal Pole** | **L** | **27** | **7.76** | **-30** | **46** | **-14** |
| **Inferior Frontal Gyrus, pars opercularis** | **L** | **17** | **7.00** | **-50** | **11** | **11** |
| **IC 52** | | | | | | |
| **Angular Gyrus** | **R** | **1860** | **22.56** | **27** | **-69** | **44** |
| Angular Gyrus | R | | 21.55 | 39 | -76 | 27 |
| Inferior Temporal Gyrus, temporooccipital part | R | | 20.54 | 52 | -59 | -12 |
| **Lateral Occipital Cortex, inferior division** | **L** | **1437** | **21.63** | **-48** | **-66** | **2** |
| Angular Gyrus | L | | 21.17 | -43 | -81 | 16 |
| Angular Gyrus | L | | 20.11 | -28 | -76 | 30 |
| **Temporal Occipital Fusiform Cortex** | **L** | **24** | **12.07** | **-28** | **-51** | **-12** |
| Temporal Fusiform Cortex, posterior division | L | | 8.92 | -30 | -41 | -12 |
| **Temporal Occipital Fusiform Cortex** | **R** | **15** | **8.49** | **37** | **-44** | **-23** |

*Note.* Results are based on one-sided t-tests and FWE-corrected $p < 0.05$ at peak level with a cluster extent threshold $k = 10$.

**Table S5.** Results for significant effects of cPPI connectivity for accuracy

| | **Accuracy** | | | | | |
|---|---|---|---|---|---|---|
| *Coefficient* | *Log-Odds* | *CI* | *p* | *Log-Odds* | *CI* | *p* |
| Intercept | 2.99 | 2.38 – 3.60 | **<0.001** | 3.19 | 2.58 – 3.80 | **<0.001** |
| DMN A+C & VAN B | -0.93 | -1.66 – -0.21 | **0.012** | | | |
| Age | -0.03 | -0.28 – 0.22 | 0.815 | 0.05 | -0.28 – 0.38 | 0.754 |
| Education | -0.12 | -0.24 – 0.01 | 0.072 | -0.16 | -0.29 – -0.02 | **0.025** |
| Motion RMSD | 1.74 | -1.46 – 4.94 | 0.286 | -2.51 | -5.39 – 0.36 | 0.086 |
| Age * DMN A+C & VAN B | 2.01 | 0.89 – 3.13 | **<0.001** | | | |
| VAN B & DAN A | | | | -0.74 | -1.91 – 0.43 | 0.213 |
| Age * VAN B & DAN A | | | | -3.43 | -5.20 – -1.65 | **<0.001** |
| **Random Effects** | | | | | | |
| $\sigma^2$ | 3.29 | | | 3.29 | | |
| $\tau_{00}$ | 0.18 $_{sub}$ | | | 0.19 $_{sub}$ | | |
| | 1.71 $_{Category}$ | | | 1.71 $_{Category}$ | | |
| ICC | 0.37 | | | 0.37 | | |
| Observations | 9837 | | | 9837 | | |
| Marginal $R^2$ / Conditional $R^2$ | 0.012 / 0.373 | | | 0.013 / 0.374 | | |

*Note*. Significant effects are marked in bold. Contrasts are sum coded. P-values were obtained via likelihood ratio tests. CI Confidence interval.

**Table S6.** Results for significant effects of cPPI connectivity for response time

| | Response time | | | | | | | | | | | |
|---|---|---|---|---|---|---|---|---|---|---|---|---|
| *Coefficient* | *Estimates* | *CI* | *p* | *Estimates* | *CI* | *p* | *Estimates* | *CI* | *p* | *Estimates* | *CI* | *p* |
| Intercept | 6.50 | 6.46 – 6.54 | **<0.001** | 6.48 | 6.44 – 6.53 | **<0.001** | 6.53 | 6.49 – 6.57 | **<0.001** | 6.52 | 6.47 – 6.56 | **<0.001** |
| DMN A+C & VAN B | 0.11 | 0.03 – 0.19 | **0.005** | | | | | | | | | |
| Age | 0.03 | 0.01 – 0.06 | **0.013** | 0.07 | 0.04 – 0.10 | **<0.001** | 0.04 | 0.02 – 0.07 | **0.002** | 0.12 | 0.09 – 0.16 | **<0.001** |
| Education | 0.00 | -0.01 – 0.02 | 0.612 | -0.00 | -0.01 – 0.01 | 0.935 | 0.00 | -0.01 – 0.01 | 0.783 | 0.00 | -0.01 – 0.01 | 0.978 |
| Motion RMSD | -0.04 | -0.38 – 0.29 | 0.814 | 0.24 | -0.01 – 0.49 | 0.061 | 0.33 | 0.08 – 0.58 | **0.009** | -0.29 | -0.56 – -0.01 | **0.042** |
| Age * DMN A+C & VAN B | 0.13 | 0.02 – 0.23 | **0.017** | | | | | | | | | |
| DMN B & DAN A | | | | -0.23 | -0.31 – -0.14 | **<0.001** | | | | | | |
| Age * DMN B & DAN A | | | | 0.56 | 0.41 – 0.72 | **<0.001** | | | | | | |
| SEM & VAN B | | | | | | | 0.02 | -0.05 – 0.09 | 0.642 | | | |
| Age * SEM & VAN B | | | | | | | -0.57 | -0.79 – -0.35 | **<0.001** | | | |
| VAN B & DAN A | | | | | | | | | | -0.48 | -0.60 – -0.35 | **<0.001** |
| Age * VAN B & DAN A | | | | | | | | | | -0.44 | -0.60 – -0.28 | **<0.001** |
| **Random Effects** | | | | | | | | | | | | |
| $\sigma^2$ | 0.11 | | | 0.11 | | | 0.11 | | | 0.11 | | |
| $\tau_{00}$ | 0.01 sub | | | 0.01 sub | | | 0.01 sub | | | 0.01 sub | | |
| | 0.00 Category | | | 0.00 Category | | | 0.00 Category | | | 0.00 Category | | |
| ICC | 0.09 | | | 0.10 | | | 0.09 | | | 0.10 | | |
| Observations | 9069 | | | 9069 | | | 9069 | | | 9069 | | |
| Marginal $R^2$ / Conditional $R^2$ | 0.011 / 0.095 | | | 0.020 / 0.118 | | | 0.020 / 0.113 | | | 0.042 / 0.138 | | |

*Note*. Significant effects are marked in bold. Contrasts are sum coded. P-values were obtained via likelihood ratio tests. CI Confidence interval.

**Table S7.** Results for linear mixed-effects model on the effect of age on brain system segregation

| | Brain system segregation | | |
|---|---|---|---|
| *Coefficient* | *Estimates* | *CI* | *P* |
| Intercept | 0.66 | 0.61 – 0.71 | **<0.001** |
| Age | -0.08 | -0.13 – -0.04 | **<0.001** |
| Motion RMSD | -0.76 | -1.19 – -0.33 | **<0.001** |
| **Random Effects** | | | |
| $\sigma^2$ | 0.00 | | |
| $\tau_{00\ sub}$ | 0.00 | | |
| ICC | 0.07 | | |
| Observations | 58 | | |
| Marginal $R^2$ / Conditional $R^2$ | 0.525 / 0.559 | | |

*Note*. Significant effects are marked in bold. Contrasts are sum coded. P-values were obtained via likelihood ratio tests. CI Confidence interval.

**Table S8.** Results for mixed-effects models on the effect of brain system segregation on accuracy and response time

| | Accuracy | | | Response time | | |
|---|---|---|---|---|---|---|
| *Coefficient* | *Log-Odds* | *CI* | *p* | *Estimates* | *CI* | *p* |
| Intercept | 2.87 | 2.15 – 3.59 | **<0.001** | 6.68 | 6.61 – 6.75 | **<0.001** |
| Global segregation | 2.62 | 0.64 – 4.60 | **0.010** | -1.34 | -1.57 – -1.11 | **<0.001** |
| Age | 0.08 | -0.18 – 0.35 | 0.548 | -0.03 | -0.05 – 0.00 | 0.078 |
| Education | -0.14 | -0.27 – -0.01 | **0.042** | 0.02 | 0.01 – 0.04 | **<0.001** |
| Motion RMSD | 0.80 | -2.14 – 3.74 | 0.599 | -1.09 | -1.43 – -0.75 | **<0.001** |
| Age * Global Segregation | -5.06 | -8.22 – -1.91 | **0.002** | 1.55 | 1.20 – 1.90 | **<0.001** |
| **Random Effects** | | | | | | |
| $\sigma^2$ | 3.29 | | | 0.11 | | |
| $\tau_{00}$ | $0.17_{sub}$ | | | $0.02_{sub}$ | | |
| | $1.71_{Category}$ | | | $0.00_{Category}$ | | |
| ICC | 0.36 | | | 0.15 | | |
| Observations | 9837 | | | 9069 | | |
| Marginal $R^2$ / Conditional $R^2$ | 0.010 / 0.370 | | | 0.052 / 0.197 | | |

Supplementary Tables

*Note*. Significant effects are marked in bold. Contrasts are sum coded. P-values were obtained via likelihood ratio tests. CI Confidence interval.

**Table S9.** Results for linear mixed-effects model on the effect of age on global efficiency

| | Global efficiency | | |
|---|---|---|---|
| *Coefficient* | *Estimates* | *CI* | *p* |
| Intercept | 0.11 | 0.10 – 0.13 | **<0.001** |
| Age | 0.02 | 0.01 – 0.03 | **<0.001** |
| Motion RMSD | -0.08 | -0.16 – -0.01 | **0.034** |
| **Random Effects** | | | |
| $\sigma^2$ | 0.00 | | |
| $\tau_{00\ sub}$ | 0.00 | | |
| ICC | 0.17 | | |
| Observations | 58 | | |
| Marginal $R^2$ / Conditional $R^2$ | 0.518 / 0.600 | | |

*Note*. Significant effects are marked in bold. Contrasts are sum coded. P-values were obtained via likelihood ratio tests. CI Confidence interval.

**Table S10.** Results for mixed-effects models on the effect of global efficiency on accuracy and response time

| | Accuracy | | | Response time | | |
|---|---|---|---|---|---|---|
| *Coefficient* | *Log-Odds* | *CI* | *p* | *Estimates* | *CI* | *p* |
| Intercept | 2.94 | 2.19 – 3.68 | **<0.001** | 6.39 | 6.34 – 6.45 | **<0.001** |
| Global efficiency | 13.62 | 4.78 – 22.45 | **0.003** | 0.66 | -0.21 – 1.53 | 0.138 |
| Age | 0.14 | -0.16 – 0.43 | 0.361 | 0.04 | 0.02 – 0.07 | **0.003** |
| Education | -0.10 | -0.23 – 0.04 | 0.157 | 0.01 | -0.00 – 0.02 | 0.239 |
| Motion RMSD | 0.94 | -1.95 – 3.82 | 0.524 | 0.63 | 0.35 – 0.91 | **<0.001** |
| Age * Global efficiency | -12.48 | -33.89 – 8.94 | 0.181 | -5.76 | -7.49 – -4.02 | **<0.001** |
| **Random Effects** | | | | | | |
| $\sigma^2$ | 3.29 | | | 0.11 | | |
| $\tau_{00}$ | 0.24 $_{sub}$ | | | 0.01 $_{sub}$ | | |
| | 1.71 $_{Category}$ | | | 0.00 $_{Category}$ | | |
| ICC | 0.37 | | | 0.10 | | |
| Observations | 9837 | | | 9069 | | |

| | | |
|---|---|---|
| Marginal $R^2$ / Conditional $R^2$ | 0.009 / 0.377 | 0.019 / 0.114 |

*Note*. Significant effects are marked in bold. Contrasts are sum coded. P-values were obtained via likelihood ratio tests. CI Confidence interval.

**Table S11.** Results for linear mixed-effects model on the effect of age on brain segregation as a function of network type

| | **Network segregation** | | |
|---|---|---|---|
| *Coefficient* | *Estimates* | *CI* | *p* |
| Intercept | 0.50 | 0.46 – 0.55 | **<0.001** |
| Age | -0.11 | -0.13 – -0.08 | **<0.001** |
| DMN A+C | 0.22 | 0.18 – 0.26 | **<0.001** |
| DMN B | 0.06 | 0.02 – 0.10 | **0.006** |
| SEM | 0.02 | -0.02 – 0.06 | 0.340 |
| CONT B | 0.13 | 0.09 – 0.17 | **<0.001** |
| VAN B | 0.17 | 0.13 – 0.21 | **<0.001** |
| DAN A | 0.31 | 0.27 – 0.35 | **<0.001** |
| Motion RMSD | -0.66 | -0.97 – -0.35 | **<0.001** |
| **Random Effects** | | | |
| $\sigma^2$ | 0.01 | | |
| $\tau_{00\ sub}$ | 0.00 | | |
| ICC | 0.10 | | |
| Observations | 406 | | |
| Marginal $R^2$ / Conditional $R^2$ | 0.551 / 0.596 | | |

*Note*. Significant effects are marked in bold. Contrasts are sum coded. P-values were obtained via likelihood ratio tests. CI Confidence interval.

**Table S12.** Results for mixed-effects models on the effect of network segregation on accuracy and response time

| | **Accuracy** | | | **Response time** | | |
|---|---|---|---|---|---|---|
| *Coefficient* | *Log-Odds* | *CI* | *p* | *Estimates* | *CI* | *p* |
| Intercept | 2.38 | 1.62 – 3.14 | **<0.001** | 6.61 | 6.49 – 6.72 | **<0.001** |
| DMN A | 0.51 | -0.24 – 1.26 | 0.183 | 0.62 | 0.52 – 0.72 | **<0.001** |

| | | | | | | |
|---|---|---|---|---|---|---|
| DMN A+C | -0.30 | -1.68 – 1.08 | 0.670 | -0.25 | -0.47 – -0.04 | **0.023** |
| DMN B | 1.07 | -0.11 – 2.25 | 0.075 | 0.23 | 0.06 – 0.40 | **0.009** |
| SEM | -0.26 | -1.20 – 0.68 | 0.590 | -0.60 | -0.73 – -0.47 | **<0.001** |
| CONT B | 1.67 | 0.07 – 3.27 | **0.040** | -0.38 | -0.64 – -0.13 | **0.003** |
| VAN B | 1.24 | 0.15 – 2.34 | **0.026** | -0.18 | -0.34 – -0.02 | **0.028** |
| DAN A | -2.22 | -3.47 – -0.96 | **0.001** | -0.32 | -0.45 – -0.18 | **<0.001** |
| Age | -0.16 | -0.52 – 0.20 | 0.390 | -0.10 | -0.16 – -0.05 | **<0.001** |
| Education | -0.11 | -0.25 – 0.04 | 0.143 | -0.02 | -0.04 – -0.01 | **0.014** |
| Motion RMSD | 4.40 | 0.93 – 7.88 | **0.013** | -0.47 | -0.98 – 0.04 | 0.073 |
| Age * DMN A | 1.30 | -0.13 – 2.73 | 0.075 | -0.84 | -1.02 – -0.65 | **<0.001** |
| Age * DMN A+C | -0.94 | -3.59 – 1.72 | 0.489 | -0.33 | -0.72 – 0.05 | 0.091 |
| Age * DMN B | -2.97 | -5.32 – -0.61 | **0.014** | 0.18 | -0.16 – 0.51 | 0.297 |
| Age * SEM | 2.18 | -0.18 – 4.54 | 0.070 | 0.39 | -0.10 – 0.89 | 0.119 |
| Age * CONT B | 1.94 | -0.76 – 4.63 | 0.159 | 0.81 | 0.47 – 1.15 | **<0.001** |
| Age * VAN B | -5.02 | -7.24 – -2.80 | **<0.001** | -0.08 | -0.40 – 0.24 | 0.607 |
| Age * DAN A | -0.19 | -2.93 – 2.55 | 0.891 | 1.68 | 1.28 – 2.07 | **<0.001** |
| **Random Effects** | | | | | | |
| $\sigma^2$ | 3.29 | | | 0.11 | | |
| $\tau_{00}$ | 0.07 $_{sub}$ | | | 0.04 $_{sub}$ | | |
| | 1.71 $_{Category}$ | | | 0.00 $_{Category}$ | | |
| ICC | 0.35 | | | 0.30 | | |
| Observations | 9837 | | | 9069 | | |
| Marginal $R^2$ / Conditional $R^2$ | 0.033 / 0.372 | | | 0.183 / 0.432 | | |

*Note*. Significant effects are marked in bold. Contrasts are sum coded. P-values were obtained via likelihood ratio tests. CI Confidence interval.

**Table S13.** Connector hubs in older adults

| Network | Region of interest | Mean PC |
|---|---|---|
| DMN A | Lateral occipital cortex, superior division 1 | 0.56 |
| DMN A | Angular gyrus 3 | 0.54 |
| DMN A+C | Lateral occipital cortex, superior division 1 | 0.56 |
| DMN A+C | Lateral occipital cortex, superior division 5 | 0.59 |
| DMN B | Angular gyrus | 0.56 |
| DMN B | Middle temporal gyrus, posterior division 2 | 0.54 |
| SEM | Paracingulate gyrus 2 | 0.56 |
| SEM | Inferior temporal gyrus, temporooccipital part | 0.53 |
| CONT B | Frontal pole 1 | 0.54 |
| CONT B | Frontal pole 3 | 0.56 |
| CONT B | Middle frontal gyrus 3 | 0.55 |
| CONT B | Angular gyrus 1 | 0.56 |
| CONT B | Angular gyrus 2 | 0.61 |
| CONT B | Superior frontal gyrus 1 | 0.54 |
| CONT B | Paracingulate gyrus | 0.57 |

*Note.* PC participation coefficient.

**Table S14.** Connector hubs in young adults

| Network | Region of interest | Mean PC |
|---|---|---|
| DMN A | Lateral occipital cortex, superior division 1 | 0.63 |
| DMN A | Angular gyrus 3 | 0.63 |
| DMN A+C | Lateral occipital cortex, superior division 1 | 0.54 |
| DMN A+C | Lateral occipital cortex, superior division 5 | 0.51 |
| DMN B | Angular gyrus | 0.58 |
| DMN B | Middle temporal gyrus, posterior division 2 | 0.52 |
| DMN B | Middle temporal gyrus, posterior division 4 | 0.51 |
| SEM | Paracingulate gyrus 2 | 0.52 |
| CONT B | Middle frontal gyrus 3 | 0.53 |
| CONT B | Angular gyrus 1 | 0.58 |
| CONT B | Angular gyrus 2 | 0.53 |
| CONT B | Lateral occipital cortex, superior division 2 | 0.58 |
| CONT B | Frontal pole 6 | 0.52 |

*Note.* PC participation coefficient.

**Table S15.** Significant age differences in participation coefficient

| Network | Region of interest | Term | Estimate | SE | t | p |
|---|---|---|---|---|---|---|
| SEM | STG, anterior division | Age | -0.16 | 0.04 | -4.28 | 0.008 |
| CONT B | Frontal pole 3 | Age | -0.13 | 0.03 | -3.92 | 0.025 |
| DAN A | Fusiform gyrus 1 | Age | -0.21 | 0.05 | -4.7 | 0.002 |
| DAN A | Fusiform gyrus 2 | Age | -0.17 | 0.04 | -4.01 | 0.02 |

*Note.* P-values are corrected for familywise error with Bonferroni-Holm method, SE standard error, STG superior temporal gyrus
